# Using deep-learning to obtain calibrated individual disease and ADL damage transition probabilities between successive ELSA waves

**DOI:** 10.1101/2023.10.24.563857

**Authors:** Emre Dil, Andrew Rutenberg

## Abstract

We predictively model damage transition probabilities for binary health outputs of 19 diseases and 25 activities of daily living states (ADLs) between successive waves of the English Longitudinal Study of Aging (ELSA). Model selection between deep neural networks (DNN), random forests, and logistic regression found that a simple one-hidden layer 128-node DNN was best able to predict future health states (AUC ≥ 0.91) and average damage probabilities (*R*^2^ ≥ 0.92). Feature selection from 134 explanatory variables found that 33 variables are sufficient to predict all disease and ADL states well. Deciles of predicted damage transition probabilities were well calibrated, but correlations between predicted health states were stronger than observed. The hazard ratios (HRs) between high-risk deciles and the average were between 3 and 10; high prevalence damage transitions typically had smaller HRs. Model predictions were good across all individual ages. A simple one-hidden layer DNN predicts multiple binary diseases and ADLs with well calibrated damage and repair transition probabilities.

## I. INTRODUCTION

Aging is the decline in healthy functioning of an organism with time [1]. Disease and disability occurrences are important discrete events during aging. Age is also a significant factor that impacts disease susceptibility, progression, and outcomes across various health conditions. The prevalence of many chronic diseases and dysfunction increase with age [2, 3]. Underlying this age dependence is a high-dimensional biological process of increasing dysfunction [4–7]. Using this high-dimensional information is important for understanding how to better predict and mitigate transitions such as disease and disability.

Using machine learning (ML) methods on high-dimensional health and medicine datasets dates is increasingly popular. ML models can perform very well on hundreds or thousands of explanatory variables simultaneously, and even for multi-dimensional target variables [8, 9]. While training models with cross-sectional data can give insights about relationships among the present health states, training with longitudinal data is needed to give insights about *future* health states [10]. Determining the “best” predictive model depends on various factors, including the nature of the data and the specific outcomes predicted. Different machine learning algorithms may be appropriate depending on the characteristics of the data and the specific predictive task [11].

We are motivated to address questions raised by three recent studies investigating aging dynamics. In the first, Farrell et. al. [12] built a computational dynamic joint interpretable network (DJIN) model to predict aging trajectories of continuous health variables from the English Longitudinal Study of Ageing (ELSA). Though comprehensive, the approach was complex and difficult to replicate – furthermore only continuous variables were predicted. We hypothesize that predictive models built around binary health-states could be significantly simpler. In the second study, Farrell et. al. [13] characterized damage and repair rates of discrete activities of daily living (ADLs) in the ELSA dataset. This study showed significant heterogeneity among damage rates, indicating the possible need for ML methods for predictive studies of binary health states. Disease transitions were not characterized. In the third study, Buergel et. al. [14] used deep-neural networks (DNN) to risk-stratify individuals for 24 common conditions, including diseases, using UK Biobank metabolomic profiles. We wanted to explore similar questions with ELSA data – without metabolomic inputs but including ADLs.

Risk stratification categorizes individuals based on their likelihood of experiencing certain events or outcomes. This facilitates tailoring interventions and treatments to different risk groups. One metric of risk stratification is the Hazard Ratio (HR), which quantifies the ratio of the hazard rates between two groups. Typically, HR values help in identifying the degree of risk association, where an HR greater than 1 signifies a higher risk in the exposed group compared to the reference group. Calibration, also important, assesses the agreement between predicted and observed risks by damage transition probabilities [15]. A well-calibrated model provides accurate risk predictions, facilitating decision-making.

ELSA [16] includes many binary, continuous, and categorical variables describing physical and mental health states, ability to do daily activities, and demographic and socioeconomic information that are repeatedly measured over different waves. As with many population studies [17], there are tens of thousands of individuals with hundreds of variables (dimensions). Our focus will be on initial waves that have both core (typically binary) and nurse (typically continuous lab) variables. We will predict subsequent individual health states by using explanatory data from the previous waves. From these health states, we will obtain predicted individual damage probabilities (i.e. individual risks).

There are two questions we want to address. The first is whether a simple deep-learning pipeline performs well for discrete health states and transitions using the available ELSA data – in particular whether it represents a significant improvement over simpler logistic-regression models (see e.g. [18, 19]) or not [20, 21]. The second is what aspects of high-dimensional input data are useful for these predictions – in particular whether discrete or continuous inputs are more useful.

## II. METHODS AND MODEL SELECTION

We consider the core and nurse data from the ELSA dataset for our study. Because the even numbered waves include the nurse data we used variables in even numbered waves as explanatory data, and health states variables in odd numbered waves as target data to predict. We consider different machine learning (ML) architectures: deep neural networks (DNN), random forest (RF) and logistic regression (LR) as predictive models. We use LR as a baseline comparison model because it is standard, interpretable, less prone to overfitting, and can sometimes surpass ML models as a clinical prediction model [20]. In contrast, DNN and RF are computationally efficient predictive models appropriate for more complex non-linear tasks – such as (we hypothesize) disease and disability during aging. We follow the study flowchart given in Fig. 1.

**FIG. 1:**
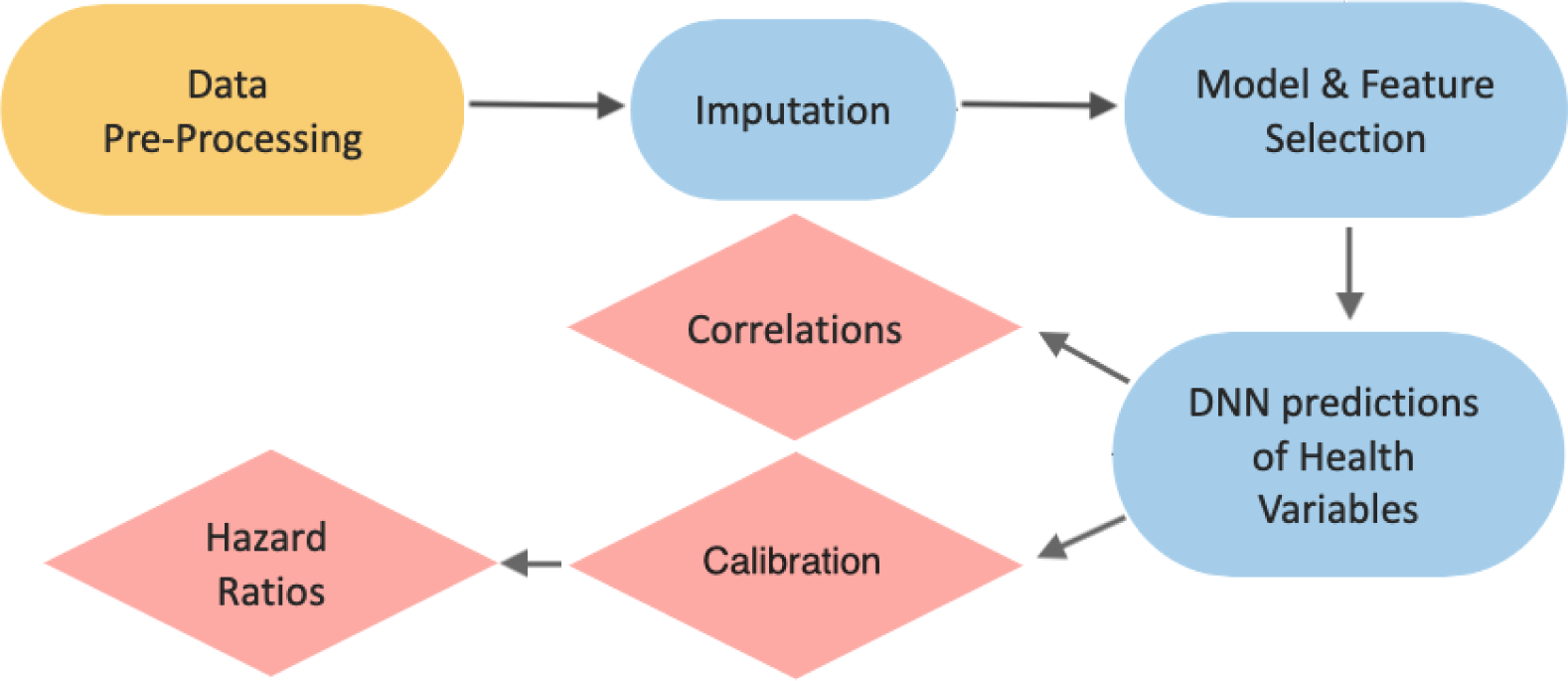
Study flowchart. In pre-processing (yellow oval), we choose target and explanatory variables, convert string variables to numbers, and normalize continuous data. We then determine the best imputation method using the MICE package in R and apply it to all data. With imputed data we apply feature and model selection to determine the best model, model architecture, and a sufficient feature-set for good predictions. Using the best model (DNN) we then predict the 19 disease and 25 ADL health states. For these predictions, we specifically explore (red diamonds) calibration, hazard ratios, and correlations.

### A. Dataset

The English Longitudinal Study of Ageing (ELSA) [16] provides information on the health dynamics and well being of an English population over 50 year old. Since 1998, there have been 9 waves separated by approximately 2 years. A summary of the dataset can be found in Table I. Waves 2, 4, 6, 8 and 9 are associated with nurse visits (“nurse” waves) where continuous lab data such as blood pressure, glucose level, or cholesterol level is collected and body measurements like height, weight, hand grip strength are taken. Data pre-processing included harmonizing variable names across waves, converting string data types to discrete numerical values, and normalizing continuous variables. We considered 134 explanatory variables with missingness [22] less than 30% from core and nurse data: 78 binary variables (including diseases and ADLs), 14 continuous variables from the core data and 42 variables mostly continuous from the nurse data. We used these 134 binary and continuous explanatory variables to predict the target health state variables in subsequent odd numbered waves. The target variables (19 diseases and 25 ADLs) are listed in Supplementary Tables S1-S2, along with their prevalences for waves 1-9, and the remaining explanatory variables are in Supplementary Table S3.

**TABLE I:**
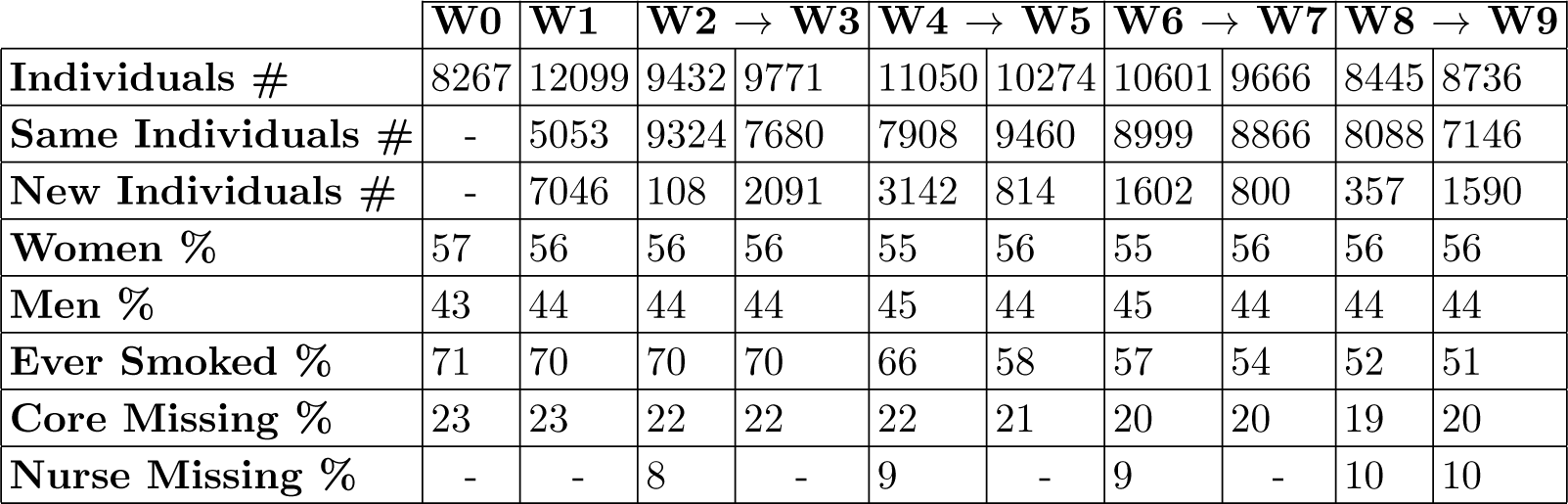
Summary of ELSA data. W*N* stands for Wave *N* of the ELSA dataset. We model four transitions that start with waves that have nurse (lab) data (waves 2, 4, 6, and 8), as indicated by the arrows. Individuals # is the number of study participants in each wave. Same individuals # is the number of participants who were also in a previous wave. New Individuals # is the number of newly added participants in the indicated wave. Core Missing % and Nurse Missing % are the missing data percentage of core data and nurse (lab) data in the corresponding wave after choosing our 134 explanatory variables. We used W0 and W1 for additional imputation input.

|  | W0 | W1 | W2 → W3 |  | W4 → W5 |  | W6 → W7 |  | W8 → W9 |  |
| --- | --- | --- | --- | --- | --- | --- | --- | --- | --- | --- |
| Individuals # | 8267 | 12099 | 9432 | 9771 | 11050 | 10274 | 10601 | 9666 | 8445 | 8736 |
| Same Individuals # | - | 5053 | 9324 | 7680 | 7908 | 9460 | 8999 | 8866 | 8088 | 7146 |
| New Individuals # | - | 7046 | 108 | 2091 | 3142 | 814 | 1602 | 800 | 357 | 1590 |
| Women % | 57 | 56 | 56 | 56 | 55 | 56 | 55 | 56 | 56 | 56 |
| Men % | 43 | 44 | 44 | 44 | 45 | 44 | 45 | 44 | 44 | 44 |
| Ever Smoked % | 71 | 70 | 70 | 70 | 66 | 58 | 57 | 54 | 52 | 51 |
| Core Missing % | 23 | 23 | 22 | 22 | 22 | 21 | 20 | 20 | 19 | 20 |
| Nurse Missing % | - | - | 8 | - | 9 | - | 9 | - | 10 | 10 |

### B. Imputation

We first perform a selection procedure to find the best imputation method and missingness mechanism for our 134 explanatory variables. We evaluate different imputation methods within the multiple imputation by chained equations (MICE) package in R version 4.2.0 (2022-04-22)[23], together with a simple reference method of imputing continuous variables by means or categorical variables by modes.

We consider both missing at random (MAR) and missing not at random (MNAR) mechanisms for MICE imputation methods. For both mechanisms, we consider both predictive mean matching (PMM) and classification and regression trees (CART) as imputation methods. We tested the best imputation method by predicting the 19 diseases’ binary health states in each wave from the imputed predictors. We compare the average accuracy, Youden’s index (*J*) and the area under curve (AUC) values for all binary predictions over all waves. Note that *AUC* ≈ (*J* + 1)/2 [24] and ranges from 0 to 1 (best). Youden’s index ranges from −1 to 1 (best).

We used simple one-hidden layer DNN and LR as the predictive model for selecting the best imputation method. By passing the imputed variables in all waves (from 0-th to 9-th waves), we predict the 19 disease states in the same wave. Then, we obtain the accuracy, Youden’s index, and AUC values for 9 waves and take their averages for both DNN and LR used as predictive models. We see in Table II for DNN and in Supplementary Table S4 for LR that MICE with MNAR is significantly better than both MICE with MAR and with the simple mean/mode matching. PMM and CART methods are quite similar. The CART method has been shown to perform better in imputation of NHANES aging data [25]. We will therefore use MICE with MNAR and the CART method to impute missing data of all input variables, and for all models.

**TABLE II:** Performance of imputation methods. Performance is evaluated with accuracy, Youden’s index, and AUC (higher is better) for DNN predictions of 19 disease states in all waves. Mean/Mode is a simple reference imputation method using mode for categorical and mean for continuous missing variables. For imputation with MICE, we considered MAR and MNAR missingness mechanisms, and PMM (default) and CART methods.

|  | Accuracy |  | Youden's J |  | AUC |  |
| --- | --- | --- | --- | --- | --- | --- |
| Mean/Mode | 0.91 |  | 0.43 |  | 0.76 |  |
|  | PMM | CART | PMM | CART | PMM | CART |
| **MICE-MAR** | 0.90 | 0.90 | 0.35 | 0.36 | 0.70 | 0.70 |
| **MICE-MNAR** | 0.93 | 0.93 | 0.53 | 0.52 | 0.80 | 0.79 |

### C. Model and Feature Selection

Our prediction models generally determine the predicted binary health variables *X_i_* (at the next wave) and its transition probability *P_i_*, for a given individual *i*. Given the binary health variables *X_i_* at the current wave, we obtain damage probability *D_i_* = (1 − *X_i_*)*P_i_* when *X_i_*= 0 and repair probability *R_i_* = *X_i_*(1 − *P_i_*) when *X_i_* = 1. To assess model and feature selection, we average damage and repair probabilities over the entire population:

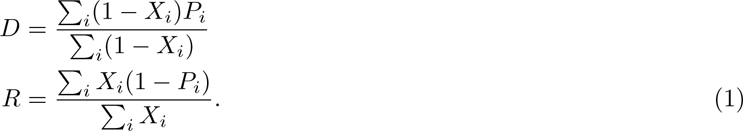

We can compare these with observed averages, in which case *P_i_* and *X_i_* are binary. For feature and model selection, we only assess the transitions from wave 2 to 3. In this paper, our focus is on damage transitions.

#### 1. Feature Selection

We performed feature selection for wave transition from 2 to 3 by choosing various number of features for each disease and ADL. We considered both filter [26] and recursive feature elimination (RFE) methods [27]. For our preferred, model agnostic, filter method, correlations or variance between features and targets are used to select features. For binary predictions, we used a f classif metric. The SelectKBest function in the Python Sklearn library was used to select a specified number of best features. The RFE method is not model agnostic and is computationally burdensome; we only considered it for the (simple) LR model.

By increasing the number of features per disease or ADL, we found that AUC rapidly saturated vs the total number of features for both DNN (see supplemental Figs. S1a and S1d) and LR (Figs. S1b and S1e) models. DNN predictions were better than LR predictions. While the AUC of disease prediction quickly plateaued at approximately 40 total features for both DNN and LR, the AUC for ADL prediction slowly increased (note small range of AUC values) as the number of features continued to increase. The performance of the RFE method with LR was worse than the filter method with LR in terms of reaching saturation very late (compare Figs. S1(b,e) and Figs. S1(c,f)). Accordingly, we used filter feature selection for all models.

The filter selected features for both disease and ADL predictions are given in Table S5 for both *N* = 33 and *N* = 41 total features. Remarkably, the same total features are selected for both disease and ADL predictions at these stages. *N* = 33 arise from selecting *k* = 2 and *k* = 9 features per disease or ADL, respectively, and dropping the duplicates. *N* = 41 arise from selecting *k* = 3 and *k* = 15, respectively. Most of the *N* = 33 features are ADL states themselves, but with cognitive diseases, pain, and grip strength as well. Added features for *N* = 41 include activity level, self-reported health, and systolic blood pressure and pulse. Significantly, very few disease states are selected – even for the goal of disease state prediction. We suspect that the small prevalence of diseases makes disease states themselves inefficient predictors. Strikingly, individual age is not selected. In Fig. S1, we observed that the reduced number of features are almost as good as full features in predictive performance for both diseases and ADLs.

#### 2. Model Selection

We considered a simple 1-hidden Layer DNN, a 2-hidden layer DNN, and an automatically generated DNN using Autokeras. In all cases, we directly predicted health states, and obtained average damage and repair probabilities using the full feature set (*N* = 134). We used binary crossentropy [28] as our loss function for DNN. We train models by using a randomly selected 80% of the total population, and use the remaining 20% as test data. We obtained the best model as the one-hidden layer DNN (1-Layer DNN) with the highest AUC (for binary health predictions) and best *R*^2^ (for average damage probabilities, by disease or ADL), see Supplementary Table S6. A 2-Hidden Layer DNN gives comparable AUC results for health state predictions. An Autokeras DNN model gave the poorest prediction results in AUC (see Table S6). Although the AUC values of health state predictions of one and two hidden layer DNNs are close to each other, the *R*^2^ values of average damage and repair probabilities estimations are slightly higher (better) for 1-Layer DNN models, see Supplementary Table S6.

We also investigated a random forest (RF) model to predict diseases and ADLs. Because prevalences are small, see Tables S1 and S2, we use the imbalanced dataset model in the imblearn [29] library in Python – we also used the Easy Ensemble Classifier with a stratified 10-Fold cross validation. For all wave transitions, our best DNN model always performs better than RF, see Supplementary Table S7. We also considered logistic regression (LR). We obtained poorer results than DNN both when predicting diseases and ADLs directly (Table S8), and when predicting damage and repair transition probabilities (Table S9). We obtained similar (worse than DNN) results even when using an LR-tuned imputation and feature selection pipeline (see Tables II and S4, and Fig. S1).

We therefore use a simple one-hidden layer DNN, as it was the best model to predict damage and repair probabilities for all wave transitions.

### D. Calibration and Hazard Ratio

We investigate the calibration of our best model using 5-fold cross validation over all waves. For a given disease or ADL we rank ordered the population with respect to predicted transition probabilities. We grouped the rank-ordered population in deciles, and compared the averages within the deciles between the model and the observed transitions. A well calibrated model has predicted and observed probabilities that are comparable [30]. A quantitative measure of calibration is the Brier score [31], which is the mean squared difference between observed and predicted probabilities. It takes values between 0 and 1, where 0 indicates the best calibration and 1 for the perfect inaccuracy in the calibration.

The hazard ratio (HR) can be used to help explain the relationship between exposures, interventions, and the occurrence of specific events. HR in our study is the ratio of the risk of the top decile over the mean risk. If all of the risk is concentrated in the top decile, then the HR will be 10. Accordingly, the HR must be between 1 and 10.

## III. RESULTS

We apply our best model, the one-hidden layer DNN with 128 nodes for the full features – for the wave transitions 2 → 3, 4 → 5, 6 → 7, and 8 → 9. The target variables are binary health states *X_i_* for the *i*th individual. The transition probabilities of health states *P_i_* can be obtained by comparing the previous and predicted future states *X_i_*, and hence average damage and repair probabilities *D* and *R* of 19 diseases and 25 ADLs can be obtained according to Eq. 1. We use AUC values to evaluate the prediction performance of binary health states, while *R*^2^ values are used to evaluate the average damage and repair probabilities of the same health variables. For diseases we find that all AUC are above 0.9 for all wave transitions, while *R*^2^ are close to 1 – see Table III. For ADLs the performance is comparable, though slightly worse. Our AUC scores are substantially better than recent deep-learning models predicting disease risk [32, 33], including a recent study by Google researchers using a large language model to analyze UK Biobank data [8]. Remarkably, our prediction performance has very little dependence on the age of individuals – see supplemental Fig. S6.

**TABLE III:** Model performance for diseases and ADLs. The evaluation performance of the best (one-layer) DNN model for full 134 features. AUC is the metric for the predictions of binary health outputs, *R*^2^ is for the average damage and repair probabilities obtained from predicted health outputs. W23 stands for the wave transition from 2 to 3, and so on.

| DISEASES |  | W23 |  | W45 |  | W67 |  | W89 |  |
| --- | --- | --- | --- | --- | --- | --- | --- | --- | --- |
|  |  | Damage | Repair | Damage | Repair | Damage | Repair | Damage | Repair |
| 1 Layer | AUC | 0.92 |  | 0.93 |  | 0.93 |  | 0.91 |  |
| | $R^2$ | 0.99 | 0.98 | 0.96 | 0.96 | 0.92 | 0.94 | 0.99 | 0.98 |

TABLE III: **Model performance for diseases and ADLs.**
| ADLs |  | W23 |  | W45 |  | W67 |  | W89 |  |
| --- | --- | --- | --- | --- | --- | --- | --- | --- | --- |
|  |  | Damage | Repair | Damage | Repair | Damage | Repair | Damage | Repair |
| 1 Layer | AUC | 0.90 |  | 0.90 |  | 0.91 |  | 0.90 |  |
| | $R^2$ | 0.94 | 0.89 | 0.97 | 0.87 | 0.97 | 0.94 | 0.96 | 0.87 |

In Fig. 2, calibration curves [15] for average of damage transition probabilities over all health states show that our model is well calibrated for both diseases and ADLs. Here, we have rank ordered predicted transition probabilities of each disease or ADL, grouped the individuals in deciles, and then averaged both predicted damage probabilities and observed damage prevalences within the decile and across diseases or ADLs. In supplemental Figs. S2 and S3 we have instead averaged the damage probabilities within individual diseases or ADLs – and also for repair probabilities. You can also see the individual disease and ADL calibration curves for damage transition probabilities of indicated waves in supplemental Figs. S4 and S5, respectively.

**FIG. 2:**
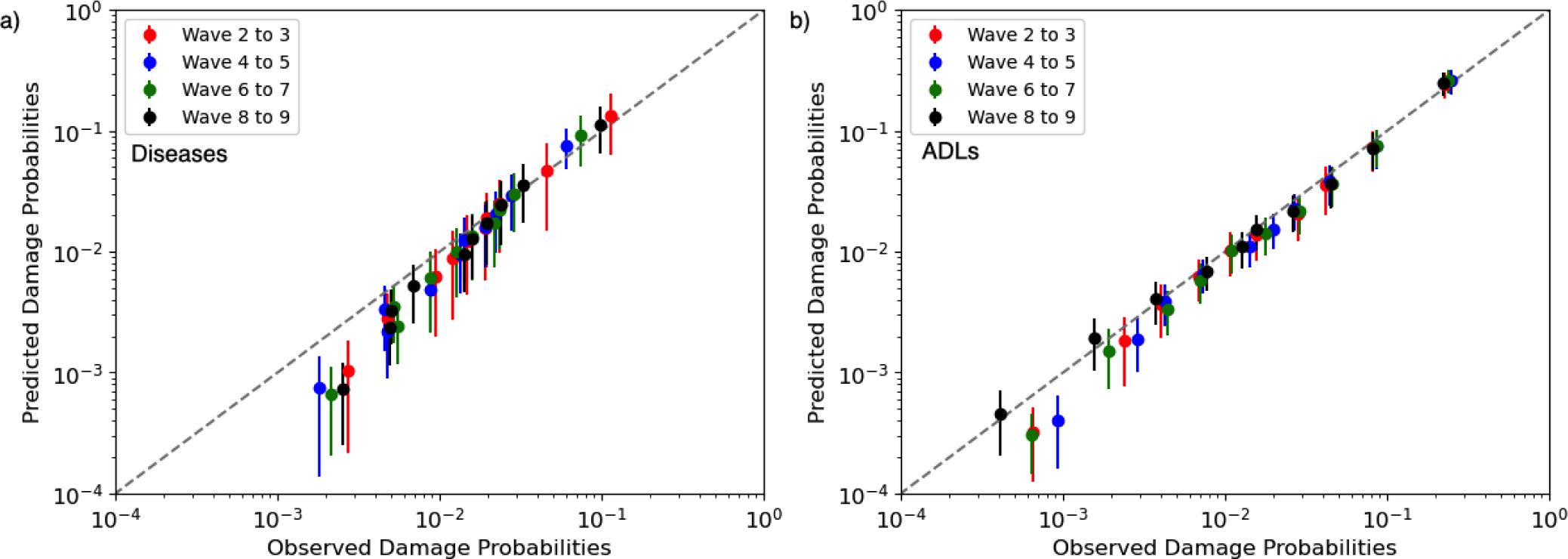
Calibration curves for damage transition probabilities. The points represent the deciles of the damage probabilities. a) For diseases, we average each decile across all diseases. See supplemental Fig. S4 for individual calibrations and supplemental Fig. S2 for averages across all deciles. b) For ADLs, we average each decile across all ADLs. See supplemental Fig. S5 for individual calibrations and supplemental Fig. S3 for averages across all deciles. Dashed line indicates perfect calibration.

We obtain a quantitative calibration measure [15] for our predictions in Supplementary Table S10 using the Brier score, which takes values between 0 and 1, with smaller values better. As seen in Table S10 the average Brier score is low for both diseases (≤ 0.020) and ADL (≤ 0.035), but diseases are on average better calibrated than ADLs. This is also reflected in the Brier scores of individual diseases and ADLs in Supplementary Tables S11 and S12.

From deciles of rank ordered damage transition probabilities, we obtain the hazard ratio (HR) values of each health variable by dividing the average probability of the maximum decile to the mean probability. These are shown in Fig. 3 for all diseases and ADLs, averaged over all wave transitions, together with standard errors from five-fold cross-validation. The dashed lines indicate the average HR. Some health conditions have low HRs, indicating that less discrimination between high and low-risk individuals was possible. Other health conditions have much higher HRs, indicating that more discrimination was possible. These results are broadly consistent across different waves (see supplemental Fig. S8). Interestingly, we find that the disease HRs are inversely correlated with disease prevalence (Spearman’s −0.43, see supplemental Fig. S7) – though the same is not seen with ADLs (Spearman’s 0.06).

**FIG. 3:**
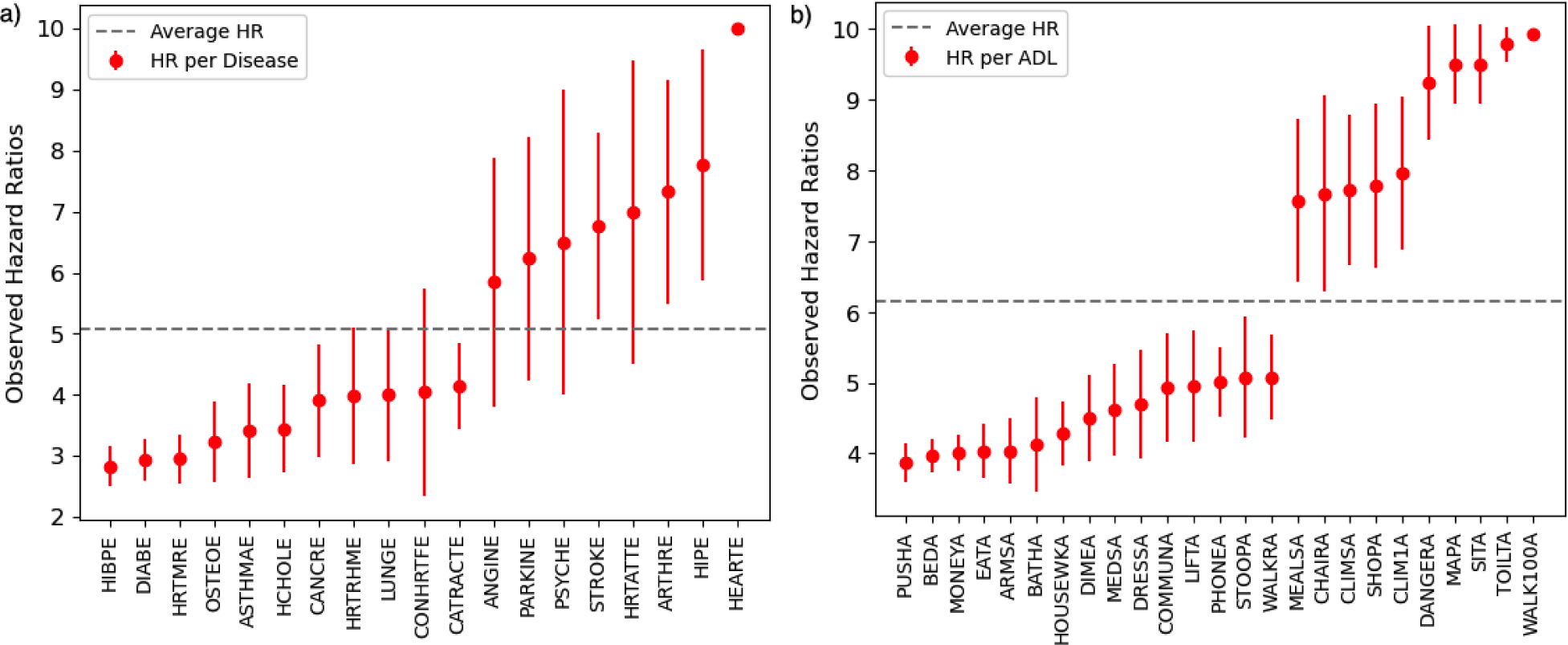
Hazard ratios. Observed hazard ratios (HR) between the highest decile of transition probability to the median for a) Diseases and b) ADLs, averaged over all wave transitions. For individual waves see Supplementary Fig. S8. Error bars are standard errors from five-fold cross-validation. The dashed line indicates the average HR.

For the wave 2 → 3 transition, we have analyzed the distribution of predicted damage transition probabilities *P_i_*’s for all diseases (see supplementary Fig. S9) and ADLs (supplementary Fig. S11). We generally find long-tailed, right-skewed distributions. With log-log plots (Figs. S10 and S12) it becomes clear that the larger probabilities are well approximated by a power-law tail with exponents typically between 2 − 3. From these distributions, we also predict bimodal character in HIBPE, DIABE and ANGINE in Fig. S9 that naturally identifies lower vs higher risk groups. We see similar high versus low risk character for HIBPE, DIABE and ANGINE from the observed and predicted HRs of high risk versus low risk group for these diseases in Supplementary Table S13 – though with HR ≤ 2.

We have also considered correlations between observed (supplementary Fig. S13) and predicted (supplementary Fig. S14) health states. We observe strong correlations between ADLs, and weaker correlations between disease states. Generally the predicted correlations are substantially stronger than observed correlations, which indicates that joint-risk is not well calibrated. However, the hierarchical clustering is similar. The DNN model may therefore be using some aspects of the correlations to improve prediction performance of individual health states and health transitions.

## IV. DISCUSSION

In this study, we predict the damage and repair probabilities for different health states using the ELSA dataset. Starting with 134 explanatory variables from a given wave, we predict future binary health states of 19 diseases and 25 ADLs in the subsequent wave. We considered several candidate models. A simple deep-neural network (DNN) with 1-hidden layer predicted future states best both with the full feature set and a selected set of 33 features (see Fig. S1(a,d), compared to other more complex DNN models, a random forest (RF) model, or a logistic regression (LR) model. Our results did not support the claim that LR is better at clinical prediction than machine learning [20]. Our best DNN always exhibited better predictive performance than LR (see Tables S8, S9, and Fig. S1). We used our best DNN model to obtain individual damage and repair probabilities for each health state. We obtained good AUC scores (approximately 0.90) for the binary health state predictions, and excellent *R*^2^ for average damage and repair probabilities.

Our selected features were largely binary health variables, with very few continuous lab/nurse variables (see supplemental Table S5 and Fig. S1). This is consistent with excellent performance when nurse data was omitted altogether (see the second section in Table S6). While initially surprising, it is consistent with the very strong correlations between health states observed in the training data (supplemental Fig. S13). Notably, while correlations among ADL variables are the strongest – the correlations between ADL and disease states are comparable to those between different disease states. We hypothesize that our DNN model has learned from this correlation structure to better predict health states – which is an advantage of developing a model that predicts all health states (disease and ADL) at once.

Our DNN damage transition probabilities for both diseases and ADLs were well calibrated (see Fig. 2 and Table S10). This means that predicted transition probabilities corresponded to observed transition probabilities when considered in rank-ordered deciles of the predicted probabilities. We used this to determine hazard-ratios (HRs) of the maximum decile to the mean. We find HRs between 3 and 10 (the largest possible for deciles) – indicating substantial risk stratification is possible with our approach. Our well-calibrated model provides accuracy in risk prediction results which would provide reliability for decision-making [34, 35]. Since we had calibrated probabilities, we also considered the distributions of transition probabilities. We found that the shape of the damage transition probability distribution was long-tailed and right-skewed – with a power-law tail with exponents typically between 2 and 3. We do not suggest any mechanism for these power-law tails.

In the feature selection of our study, age was not selected as one of the best predictors during feature selection (see Table. S5. While age is a significant risk factor for most diseases and ADLs [2, 3], the disease and ADL context indirectly provide sufficient information about that risk. Remarkably, our predictive quality does not strongly depend on age (see Fig. S6). The age at disease onset may nevertheless influence disease trajectory and long-term outcomes. Our approach has limitations. We trained each wave transition separately. We have not validated with other datasets. While damage transition probabilities were well calibrated, the repair transition probabilities were not (hence we have postponed extensive discussion of them) – and neither were the correlations between health states. It is also important to note that we predicted the disease and ADL states as reported within the data. The connection between these observed health states and the onset of the “real” underlying diseases or health conditions could not be explored.

We investigated how much information about the future health states can be found in current states, and what sort of models are best designed to make the prediction. We found that the simple one-hidden layer DNN worked best, which is encouraging for development of interpretable and useful (i.e. translational) ML models in the future. Our model is simpler than some competing ML approaches, but seems to perform as well or better. It will be interesting to explore whether our simple approach can be substantially improved, for example by using information from multiple previous waves. It will also be important to explore whether readily available clinical information can lead to sufficiently good predictions to be useful. We have shown that the most significant features were generally the ADL health states together with cognitive function, pain level and grip strength.

## ACKNOWLEDGEMENTS

ADR thanks the Natural Sciences and Engineering Research Council (NSERC) for an operating Grant (RGPIN 2019-05888).

## VI. SUPPLEMENTARY MATERIAL

Tables S1-S2: Percentage prevalence of Diseases and ADLs

Table S3: Used core and nurse variables with their code and real name.

Table S4: Performance of imputation methods based on LR predictions

Table S5: Selected features in feature selection.

Table S6: Evaluation of best DNN model selection.

Table S7: Comparison of the performances of RF and the best DNN model.

Tables S8-S9: Comparison of the performances of LR and the best DNN model for health states and transition probabilities (2 times target variables), respectively.

Table S10: Calibration scores for the average of all diseases and ADLs.

Tables S11-S12: Individual disease and ADL calibration scores.

Table S13: Observed and predicted HRs of high versus low risk groups for the bimodal distributions in Fig. S9.

Figure S1: AUC vs #n features for filter DNN, filter LR and RFE LR feature selection methods.

Figures S2-S3: Scatter plot for disease and ADL damage repair probabilities.

Figures S4-S5: Individual disease and ADL calibration curves for damage transition probabilities.

Figure S6: Prediction performances of diseases and ADLs according to age.

Figure S7: Hazard ratio versus prevalence of diseases and ADLs.

Figure S8: HR versus all diseases and ADLs over all individual waves.

Figures S9-S10: Histograms of predicted damage transition probabilities for each disease in log-lin and log-log scales.

Figures S11-S12: Histograms of predicted damage transition probabilities for each ADL in log-lin and log-log scales.

Figures S13-S14: Correlation map between observed and predicted health states.

**TABLE S1:** Percentage prevalence of 19 disease outputs: W*N* represents Wave *N* in ELSA data. Prevalences are given for waves 1-9 together with the average prevalence over all waves. Diseases have been rank ordered in increasing average prevalence.

|  | Health Output | W1 | W2 | W3 | W4 | W5 | W6 | W7 | W8 | W9 | Average |
| --- | --- | --- | --- | --- | --- | --- | --- | --- | --- | --- | --- |
| **PARKINE** | ever had Parkinson's disease | 0 | 1 | 1 | 1 | 1 | 1 | 1 | 1 | 1 | 1 |
| **CONHRTFE** | ever had congestive heart failure | 1 | 1 | 1 | 1 | 1 | 1 | 1 | 1 | 1 | 1 |
| **HIPE** | ever had hip fracture | 2 | 1 | 1 | 1 | 1 | 1 | 1 | 1 | 1 | 1 |
| **HEARTE** | ever had heart problems | 0 | 1 | 2 | 3 | 3 | 4 | 2 | 2 | 6 | 2 |
| **HRTMRE** | ever had heart murmur | 5 | 4 | 5 | 4 | 4 | 4 | 4 | 4 | 3 | 4 |
| **HRTATTE** | ever had heart attack | 0 | 6 | 6 | 6 | 6 | 5 | 5 | 5 | 1 | 4 |
| **STROKE** | ever had stroke | 4 | 5 | 5 | 4 | 5 | 5 | 5 | 5 | 3 | 5 |
| **LUNGE** | ever had lung disease | 6 | 7 | 5 | 5 | 5 | 5 | 5 | 5 | 5 | 5 |
| **CANCRE** | ever had cancer | 6 | 7 | 5 | 5 | 6 | 6 | 6 | 7 | 8 | 6 |
| **ANGINE** | ever had angina | 9 | 8 | 8 | 7 | 9 | 6 | 3 | 3 | 1 | 6 |
| **OSTEOE** | ever had osteoporosis | 5 | 6 | 6 | 7 | 8 | 7 | 8 | 8 | 8 | 7 |
| **HRTRHME** | ever had abnormal heart rhythm | 6 | 7 | 8 | 7 | 8 | 8 | 9 | 10 | 8 | 8 |
| **PSYCHE** | ever had psych problems | 8 | 9 | 8 | 9 | 10 | 10 | 10 | 10 | 8 | 9 |
| **DIABE** | ever had diabetes | 7 | 6 | 9 | 9 | 11 | 11 | 12 | 13 | 12 | 10 |
| **ASTHMAE** | ever had asthma | 12 | 13 | 11 | 11 | 11 | 11 | 11 | 11 | 11 | 11 |
| **CATRACTE** | ever had cataracts | 13 | 17 | 17 | 17 | 20 | 22 | 25 | 30 | 30 | 21 |
| **HCHOLE** | ever had high cholesterol | 0 | 18 | 29 | 32 | 36 | 34 | 35 | 36 | 32 | 28 |
| **ARTHRE** | ever had arthritis | 32 | 36 | 35 | 34 | 37 | 37 | 37 | 40 | 38 | 36 |
| **HIBPE** | ever had high blood pressure | 37 | 32 | 40 | 38 | 40 | 38 | 38 | 39 | 36 | 37 |

**TABLE S2:** Percentage prevalence of 25 ADL outputs: W*N* represents Wave *N* in ELSA data. Prevalences are given for waves 1-9 together with the Average prevalence over all waves. ADLs have been rank ordered in increasing average prevalence. ‘Diff’ = ‘Difficulty’.

|  | Health Output | W1 | W2 | W3 | W4 | W5 | W6 | W7 | W8 | W9 | Average |
| --- | --- | --- | --- | --- | --- | --- | --- | --- | --- | --- | --- |
| **DANGERA** | Some Diff-Recognizing when in physical danger | 0 | 0 | 0 | 2 | 2 | 2 | 2 | 2 | 2 | 1 |
| **EATA** | Some Diff-Eating | 2 | 2 | 2 | 2 | 2 | 3 | 2 | 3 | 3 | 2 |
| **MEDSA** | Some Diff-Take medications | 1 | 2 | 2 | 2 | 3 | 3 | 3 | 3 | 3 | 3 |
| **COMMUNA** | Some Diff-Communication (speech, hearing, eye) | 0 | 0 | 0 | 4 | 4 | 4 | 4 | 4 | 4 | 3 |
| **PHONEA** | Some Diff-Use a telephone | 2 | 2 | 3 | 3 | 3 | 3 | 3 | 3 | 3 | 3 |
| **MONEYA** | Some Diff-Managing money | 2 | 3 | 4 | 3 | 4 | 4 | 4 | 4 | 4 | 4 |
| **WALKRA** | Some Diff-Walk across room | 3 | 4 | 3 | 3 | 3 | 4 | 4 | 4 | 4 | 4 |
| **TOILTA** | Some Diff-Using the toilet | 3 | 3 | 4 | 3 | 4 | 4 | 4 | 4 | 4 | 4 |
| **MEALSA** | Some Diff-Prepare hot meal | 4 | 5 | 5 | 5 | 5 | 5 | 5 | 6 | 6 | 5 |
| **MAPA** | Some Diff-Use a map | 5 | 6 | 5 | 5 | 5 | 5 | 5 | 5 | 5 | 5 |
| **BEDA** | Some Diff-Get in/out bed | 2 | 6 | 6 | 6 | 6 | 6 | 6 | 6 | 6 | 6 |
| **DIMEA** | Some Diff-Pick up a 5p coin | 5 | 5 | 5 | 5 | 6 | 6 | 6 | 6 | 6 | 6 |
| **SHOPA** | Some Diff-Shop for grocery | 9 | 10 | 10 | 9 | 10 | 9 | 9 | 10 | 9 | 9 |
| **BATHA** | Some Diff-Bathing, shower | 12 | 12 | 11 | 10 | 10 | 10 | 9 | 9 | 9 | 10 |
| **ARMSA** | Some Diff-Rch/xtnd arms up | 11 | 11 | 11 | 10 | 11 | 12 | 11 | 11 | 10 | 11 |
| **DRESSA** | Some Diff-Dressing | 13 | 14 | 13 | 13 | 13 | 13 | 12 | 13 | 13 | 13 |
| **WALK100A** | Some Diff-Walk 100y | 12 | 12 | 12 | 12 | 14 | 14 | 13 | 14 | 13 | 13 |
| **SITA** | Some Diff-Sit for 2 hours | 14 | 14 | 14 | 12 | 13 | 13 | 13 | 13 | 11 | 13 |
| **CLIM1A** | Some Diff-Clmb 1 ft str | 15 | 15 | 14 | 14 | 15 | 15 | 15 | 15 | 14 | 15 |
| **HOUSEWKA** | Some Diff-Doing work around the house or garden | 16 | 17 | 15 | 15 | 16 | 15 | 15 | 16 | 15 | 15 |
| **PUSHA** | Some Diff-Push/pull lg obj | 18 | 19 | 18 | 17 | 18 | 18 | 17 | 18 | 16 | 18 |
| **LIFTA** | Some Diff-Lift/carry 10lbs | 26 | 25 | 24 | 23 | 23 | 23 | 22 | 23 | 21 | 23 |
| **CHAIRA** | Some Diff-Get up fr chair | 26 | 27 | 25 | 25 | 25 | 24 | 24 | 24 | 23 | 25 |
| **CLIMSA** | Some Diff-Clmb sev ft str | 36 | 38 | 34 | 34 | 35 | 32 | 31 | 33 | 30 | 34 |
| **STOOPA** | Some Diff-Stoop/Kneel/Crch | 35 | 38 | 35 | 35 | 38 | 37 | 37 | 39 | 37 | 37 |

**TABLE S3:**
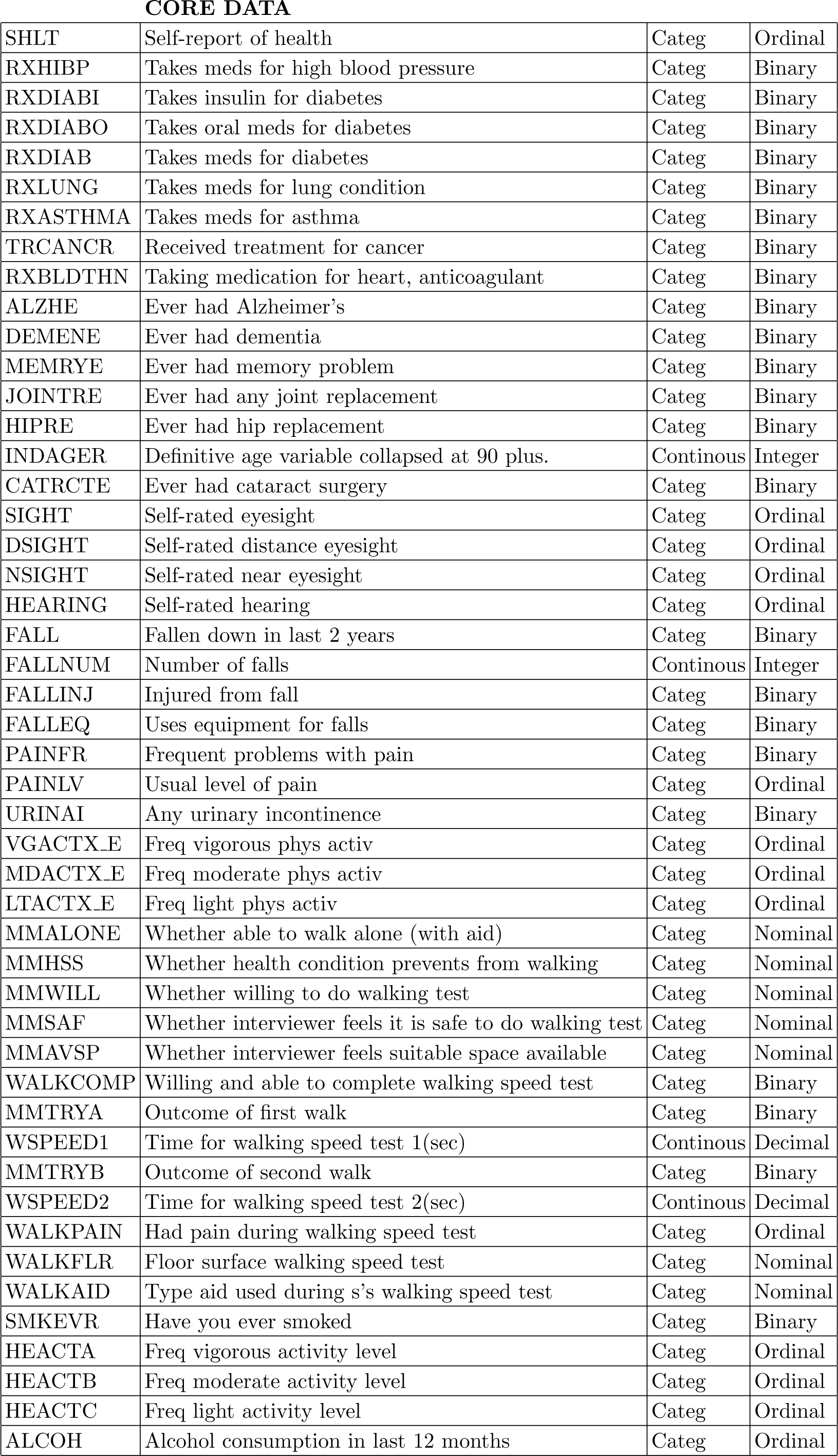
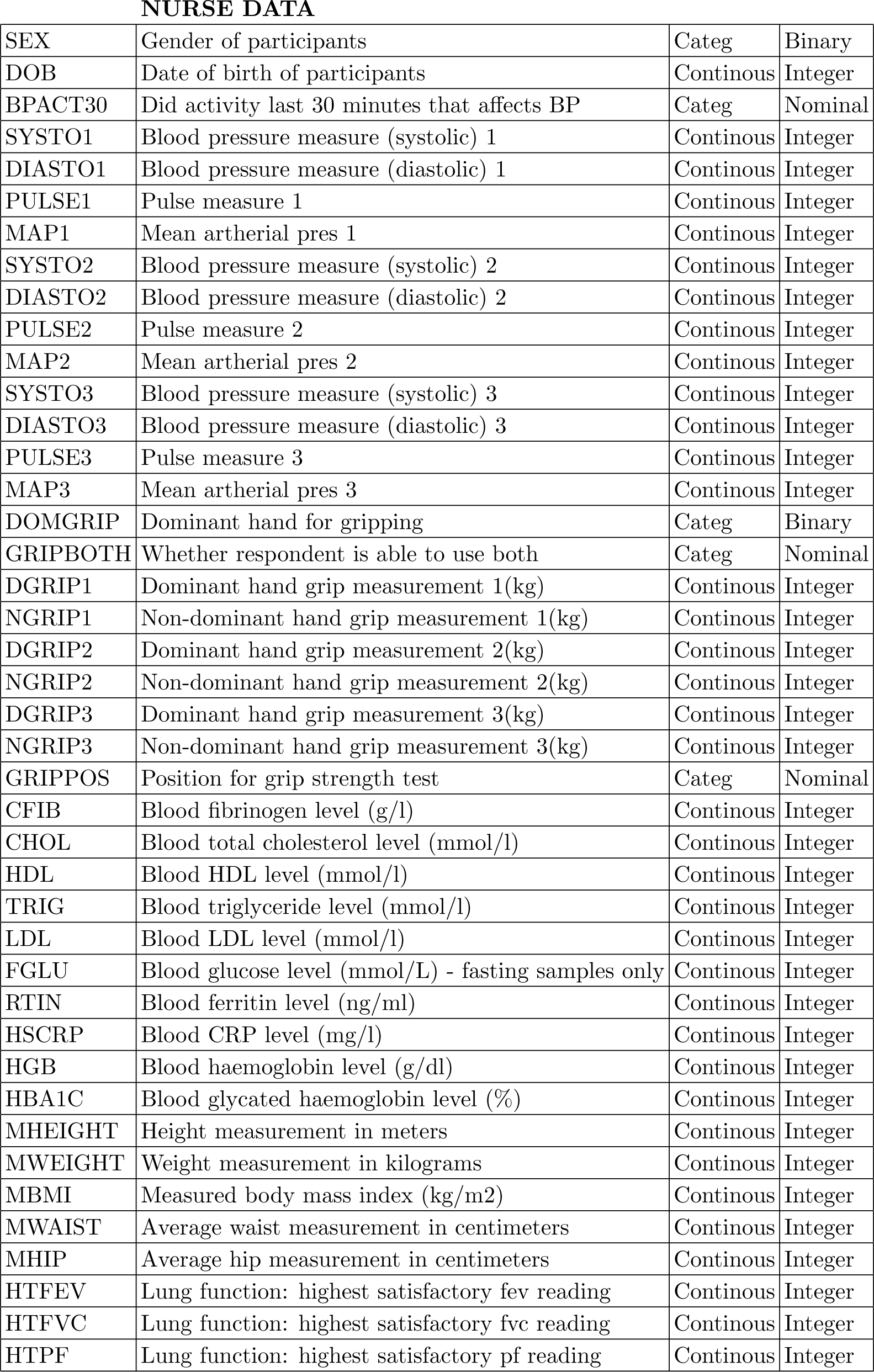
Core and Nurse Variables used in this study.

**TABLE S4:** Performance of imputation methods for LR predictions. Performance is evaluated with accuracy, Youden’s index, and AUC (higher is better). Mean/Mode is a simple reference imputation method using mode for categorical and mean for continuous missing variables. For imputation with MICE, we considered MAR and MNAR missingness mechanisms, and PMM (default) and CART methods.

|  | **Accuracy** |  | **Youden's J** |  | **AUC** |  |
| --- | --- | --- | --- | --- | --- | --- |
| Mean/Mode | 0.91 |  | 0.14 |  | 0.50 |  |
|  | **PMM** | **CART** | **PMM** | **CART** | **PMM** | **CART** |
| **MICE-MAR** | 0.90 | 0.90 | 0.05 | 0.04 | 0.50 | 0.50 |
| **MICE-MNAR** | 0.93 | 0.93 | 0.28 | 0.27 | 0.50 | 0.50 |

**TABLE S5:** Selected features using the filter feature selection method, for *N* = 33 and *N* = 41 total features (reading from the top) for both disease and ADL predictions. The variable types are also indicated. The number of iterations (*k*) indicates how many variables were chosen for each output; *N* represents the total number of unique variables for all outputs.

| Selected Features | Number of Iterations for Disease&ADL | Total Selected Features | Variable Type |
| --- | --- | --- | --- |
| WALKRA |  |  | ADL-Core Binary |
| DRESSA |  |  | ADL-Core Binary |
| BATHA |  |  | ADL-Core Binary |
| EATA |  |  | ADL-Core Binary |
| BEDA |  |  | ADL-Core Binary |
| TOILTA |  |  | ADL-Core Binary |
| MAPA |  |  | ADL-Core Binary |
| MEALSA |  |  | ADL-Core Binary |
| SHOPA |  |  | ADL-Core Binary |
| PHONEA |  |  | ADL-Core Binary |
| MEDSA |  |  | ADL-Core Binary |
| HOUSEWKA |  |  | ADL-Core Binary |
| MONEYA |  |  | ADL-Core Binary |
| WALK100A |  |  | ADL-Core Binary |
| SITA |  |  | ADL-Core Binary |
| CHAIRA |  |  | ADL-Core Binary |
| CLIMSA |  |  | ADL-Core Binary |
| CLIM1A |  |  | ADL-Core Binary |
| STOOPA |  |  | ADL-Core Binary |
| LIFTA |  |  | ADL-Core Binary |
| DIMEA |  |  | ADL-Core Binary |
| ARMSA |  |  | ADL-Core Binary |
| PUSHA |  |  | ADL-Core Binary |
| ARTHRE |  |  | Disease-Core Binary |
| ALZHE |  |  | Core Binary |
| DEMENE |  |  | Core Binary |
| MEMRYE |  |  | Core Binary |
| FALLEQ |  |  | Core Binary |
| PAINFR |  |  | Core Binary |
| HEACTB |  |  | Core Cont. |
| PAINLV |  |  | Core Cont. |
| GRIPBOTH |  |  | Nurse |
| GRIPPOS | k=2 & 9 | N=33 | Nurse |
| HEACTC |  |  | Core Cont. |
| SHLT |  |  | Core Cont. |
| INDAGER |  |  | Core Cont. |
| MDACTX_E |  |  | Core Cont. |
| SYSTO1 |  |  | Nurse |
| PULSE1 |  |  | Nurse |
| SYSTO2 |  |  | Nurse |
| SYSTO3 | k=3 & 15 | N=41 | Nurse |

**TABLE S6:**
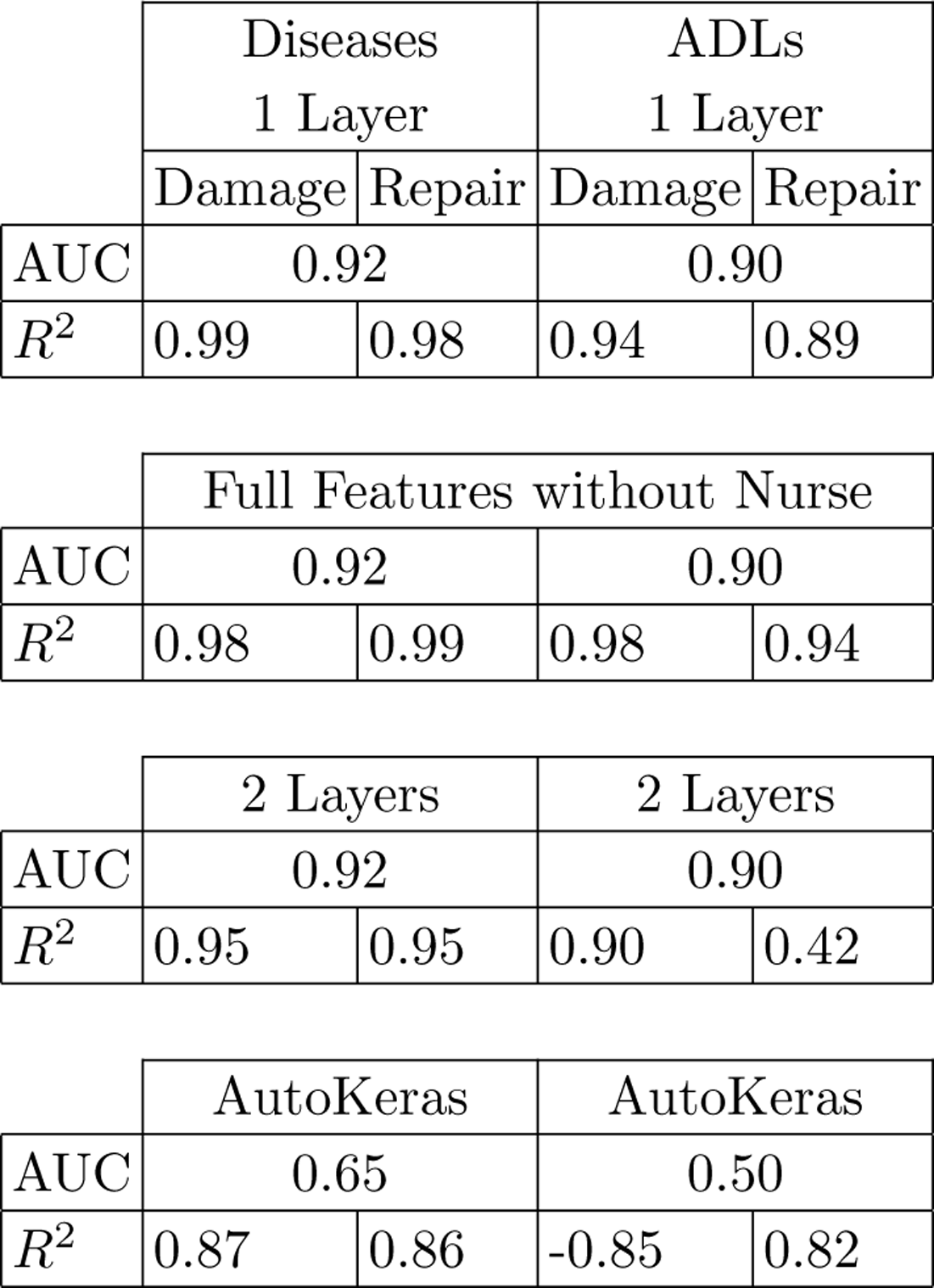
Model selection. Area under the ROC curve (AUC) is the evaluation metric for binary health state predictions, while *R*^2^ is the metric for predicted and observed (real) values of average damage and repair probabilities. The full feature set of 134 features are used for the 1 layer, 2 layer, and AutoKeras models. In terms of AUC the 1 and 2 layer models are comparable, but the AutoKeras model does not perform as well. Negative *R*^2^ shows a worse fit than the horizontal line defined by the mean of the data points. In terms of the *R*^2^ the one layer model is best, and the performance is only slightly degraded without the nurse data.

**TABLE S7:**
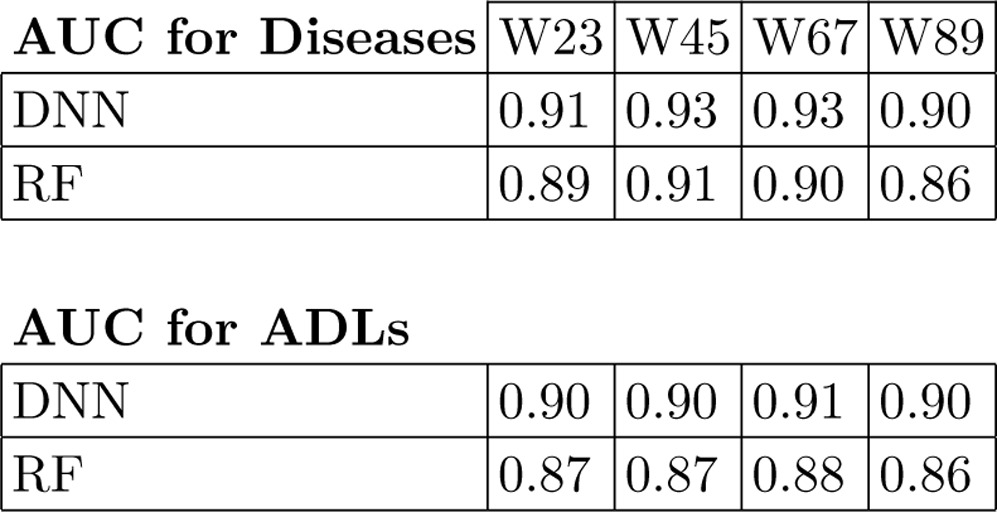
Comparison of random forest (RF) and the best DNN model for different waves, as indicated. Performance is assessed by the average AUC of disease and ADL states. The full set of 134 features were used. We see that our DNN model reliably outperforms RF.

**TABLE S8:** Comparison of the prediction performance of logistic regression (LR) and the best DNN model for 19 disease states and 25 ADLs. DNN is better than LR for both diseases and ADLs. The trivial model assumes that the target variables do not change from the previous wave. We used the full 134 features.

| DISEASES | DNN | LR | Trivial |
| --- | --- | --- | --- |
| AUC | 0.92 | 0.79 | 0.80 |

TABLE S8: Comparison of the prediction performance of logistic regression (LR) and the best DNN model for 19 disease states and 25 ADLs.
| ADLs | DNN | LR | Trivial |
| --- | --- | --- | --- |
| AUC | 0.90 | 0.69 | 0.72 |

**TABLE S9:**
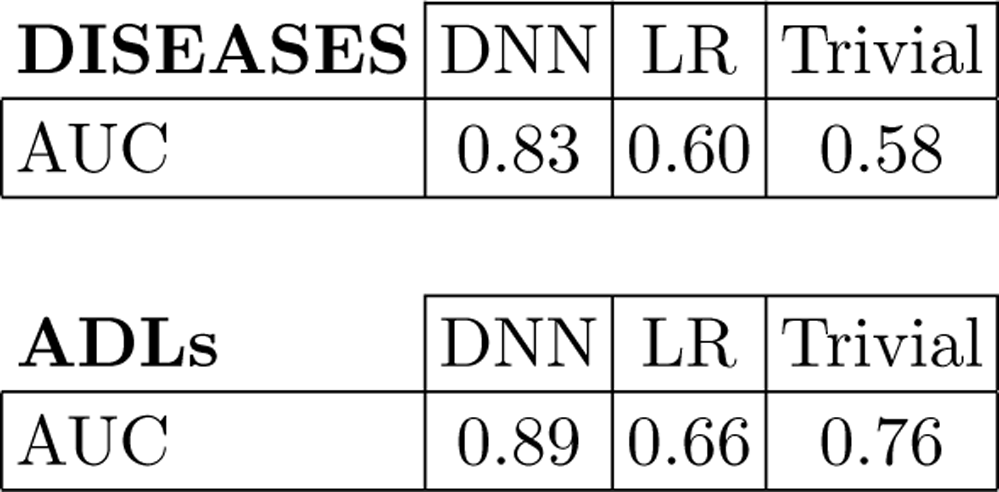
Comparison of the prediction performance of logistic regression (LR) and the best DNN model for predicting transitions of 19 disease states and 25 ADLs. DNN is better than LR for both diseases and ADLs. The trivial model assumes that the target variables all change to their opposite values from the previous wave values. We used the full 134 features.

**TABLE S10:** Calibration scores: Brier score takes values between 0 and 1, which represent the complete calibration and complete non-calibration. We average Brier scores over all diseases and ADLs and have well calibration scores for all diseases and ADLs for all wave transitions. See Tables S11 and S12 for individual disease and ADL Brier scores.

| **Brier Score** | **DISEASES** | **ADLs** |
| --- | --- | --- |
| **Wave 2 to 3** | 0.020 | 0.032 |
| **Wave 4 to 5** | 0.017 | 0.034 |
| **Wave 6 to 7** | 0.018 | 0.034 |
| **Wave 8 to 9** | 0.019 | 0.032 |

**TABLE S11:** For individual disease calibration scores: All individual diseases over all waves show a well calibration since they all have Brier scores close to 0.

| **Brier** | **Wave 2 to 3** | **Wave 4 to 5** | **Wave 6 to 7** | **Wave 8 to 9** |
| --- | --- | --- | --- | --- |
| HIBPE | 0.029 | 0.026 | 0.026 | 0.040 |
| DIABE | 0.037 | 0.026 | 0.021 | 0.025 |
| CANCRE | 0.026 | 0.022 | 0.020 | 0.022 |
| LUNGE | 0.019 | 0.015 | 0.016 | 0.016 |
| HEARTE | 0.006 | 0.005 | 0.007 | 0.007 |
| STROKE | 0.016 | 0.018 | 0.022 | 0.025 |
| PSYCHE | 0.004 | 0.003 | 0.006 | 0.002 |
| ARTHRE | 0.020 | 0.021 | 0.017 | 0.022 |
| ASTHMAE | 0.013 | 0.015 | 0.016 | 0.023 |
| HCHOLE | 0.046 | 0.034 | 0.039 | 0.036 |
| CATRACTE | 0.016 | 0.010 | 0.009 | 0.008 |
| PARKINE | 0.018 | 0.015 | 0.019 | 0.019 |
| HIPE | 0.012 | 0.005 | 0.011 | 0.010 |
| ANGINE | 0.008 | 0.015 | 0.016 | 0.017 |
| HRTATTE | 0.004 | 0.005 | 0.008 | 0.009 |
| CONHRTFE | 0.022 | 0.021 | 0.023 | 0.021 |
| HRTMRE | 0.016 | 0.017 | 0.015 | 0.027 |
| HRTRHME | 0.039 | 0.031 | 0.028 | 0.029 |
| OSTEOE | 0.021 | 0.014 | 0.016 | 0.012 |

**TABLE S12:** For individual ADL calibration scores: All individual ADLs over all waves show a well calibration since they all have Brier scores close to 0.

| **Brier** | **Wave 2 to 3** | **Wave 4 to 5** | **Wave 6 to 7** | **Wave 8 to 9** |
| --- | --- | --- | --- | --- |
| WALKRA | 0.033 | 0.032 | 0.032 | 0.034 |
| DRESSA | 0.023 | 0.027 | 0.023 | 0.026 |
| BATHA | 0.021 | 0.030 | 0.025 | 0.029 |
| EATA | 0.064 | 0.067 | 0.065 | 0.065 |
| BEDA | 0.060 | 0.062 | 0.056 | 0.066 |
| TOILTA | 0.012 | 0.014 | 0.015 | 0.013 |
| MAPA | 0.008 | 0.008 | 0.009 | 0.009 |
| DANGERA | 0.008 | 0.011 | 0.010 | 0.011 |
| MEALSA | 0.040 | 0.037 | 0.038 | 0.037 |
| SHOPA | 0.035 | 0.036 | 0.039 | 0.036 |
| PHONEA | 0.034 | 0.032 | 0.037 | 0.032 |
| COMMUNA | 0.022 | 0.025 | 0.030 | 0.022 |
| MEDSA | 0.021 | 0.027 | 0.031 | 0.026 |
| HOUSEWKA | 0.064 | 0.062 | 0.064 | 0.057 |
| MONEYA | 0.056 | 0.063 | 0.061 | 0.055 |
| WALK100A | 0.016 | 0.013 | 0.012 | 0.013 |
| SITA | 0.009 | 0.008 | 0.010 | 0.008 |
| CHAIRA | 0.009 | 0.010 | 0.010 | 0.008 |
| CLIMSA | 0.037 | 0.033 | 0.038 | 0.034 |
| CLIM1A | 0.038 | 0.033 | 0.038 | 0.034 |
| STOOPA | 0.027 | 0.037 | 0.030 | 0.027 |
| LIFTA | 0.024 | 0.028 | 0.024 | 0.022 |
| DIMEA | 0.021 | 0.031 | 0.029 | 0.027 |
| ARMSA | 0.064 | 0.068 | 0.061 | 0.061 |
| PUSHA | 0.056 | 0.060 | 0.059 | 0.056 |

**TABLE S13:** Observed and predicted HRs of high versus low risk groups for the bimodal distributions in Fig. S9.

|  | **HR of High-Low**<br>**Risk Groups** |  |
| --- | --- | --- |
|  | **Observed** | **Predicted** |
| **HIBPE** | 1.74 | 1.72 |
| **DIABE** | 1.31 | 1.38 |
| **ANGINE** | 1.94 | 1.87 |

**FIG. S1:**
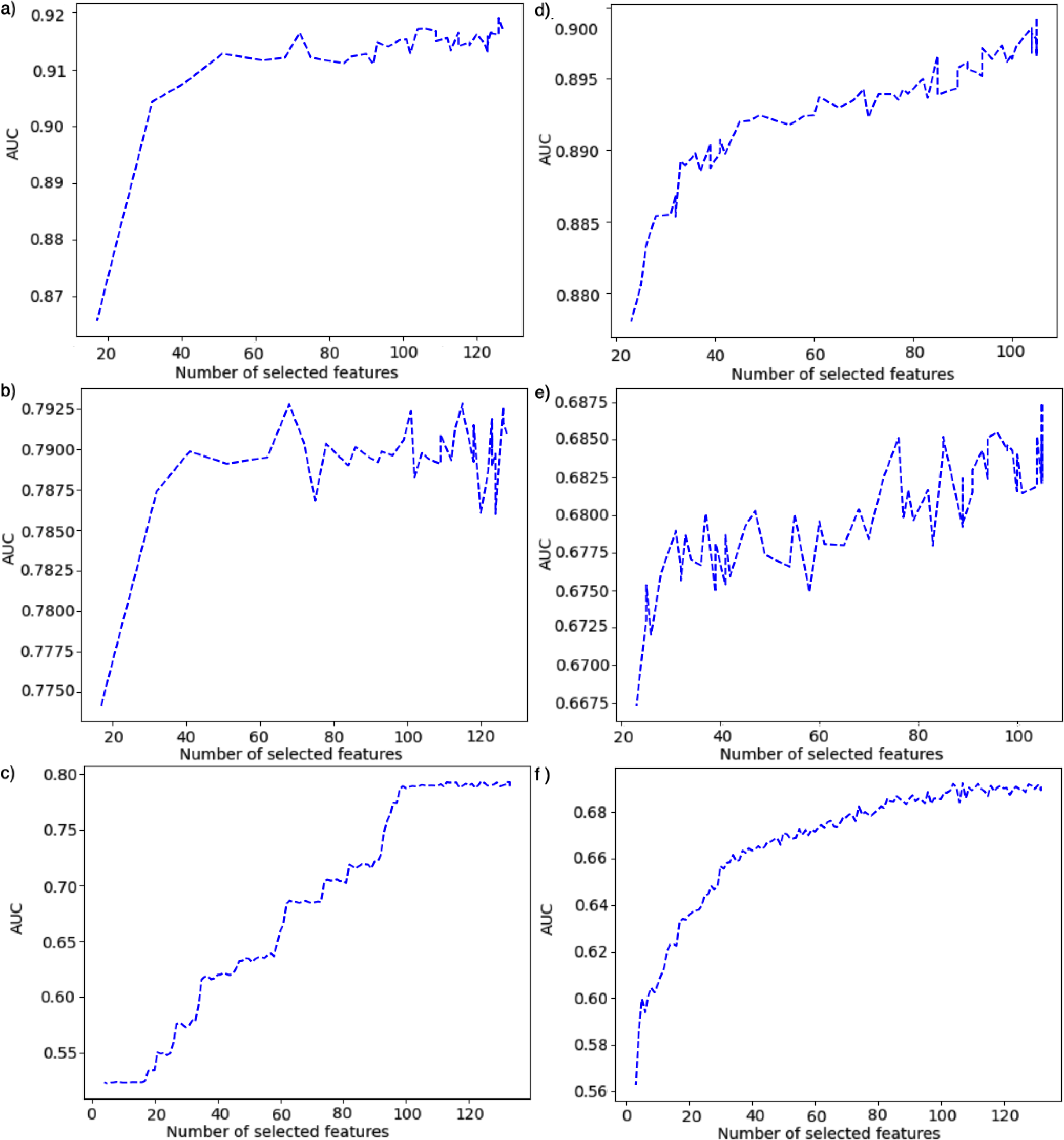
Prediction performance in terms of AUC vs #n features for diseases a) in filter DNN, b) in filter LR, c) in RFE LR methods and for ADLs d) in filter DNN, e) in filter LR, and f) in RFE LR methods. While DNN is the best model, filter method is the best feature selection method with the earliest AUC saturation for both diseases and ADLs in a) and d), respectively.

**FIG. S2:**
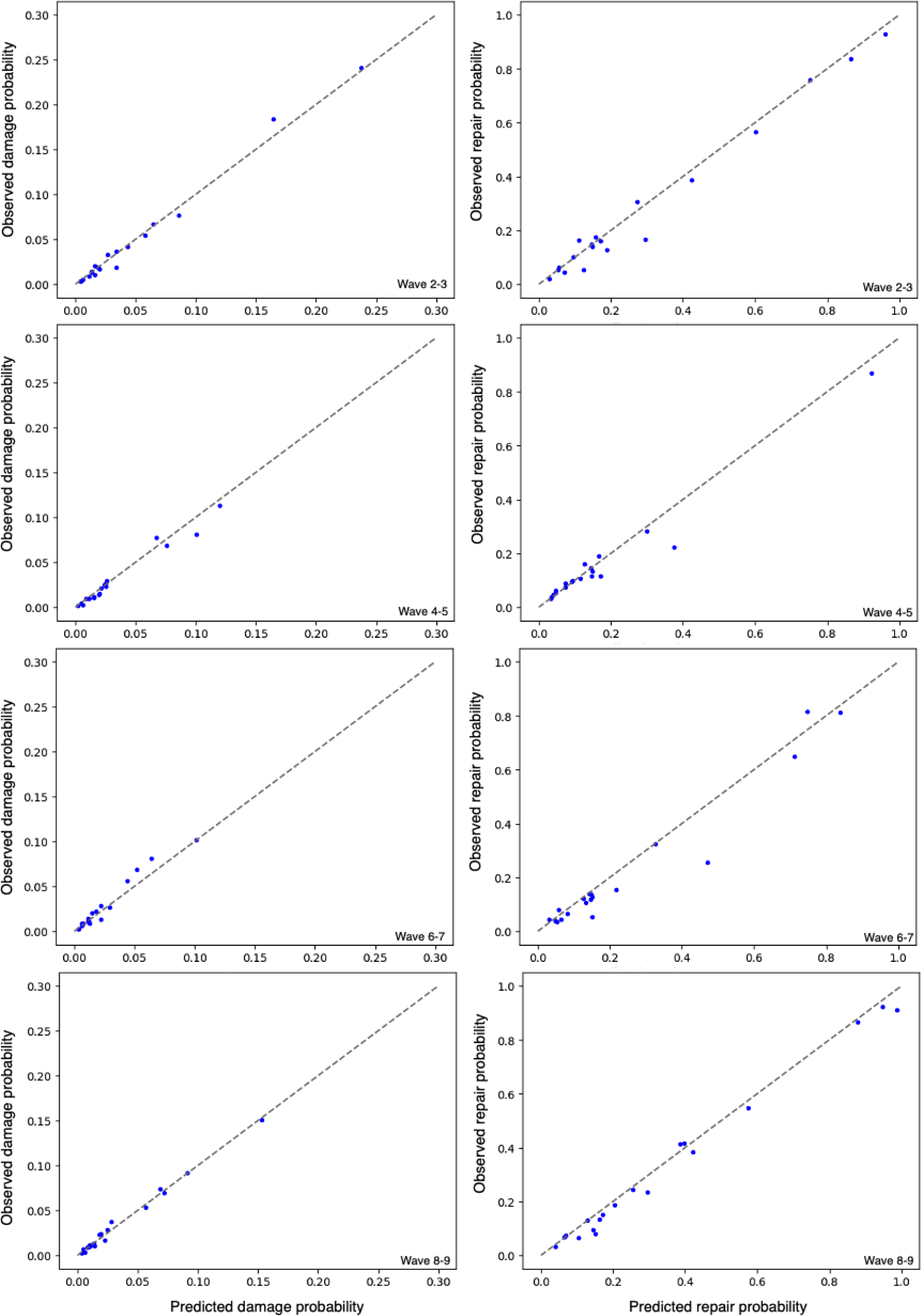
Predicted versus observed average damage and repair probabilities for Diseases. Each point represents one disease. Waves are as indicated. The average *R*^2^ across waves are in Table III Diseases. The gray dashed line indicates perfect calibration.

**FIG. S3:**
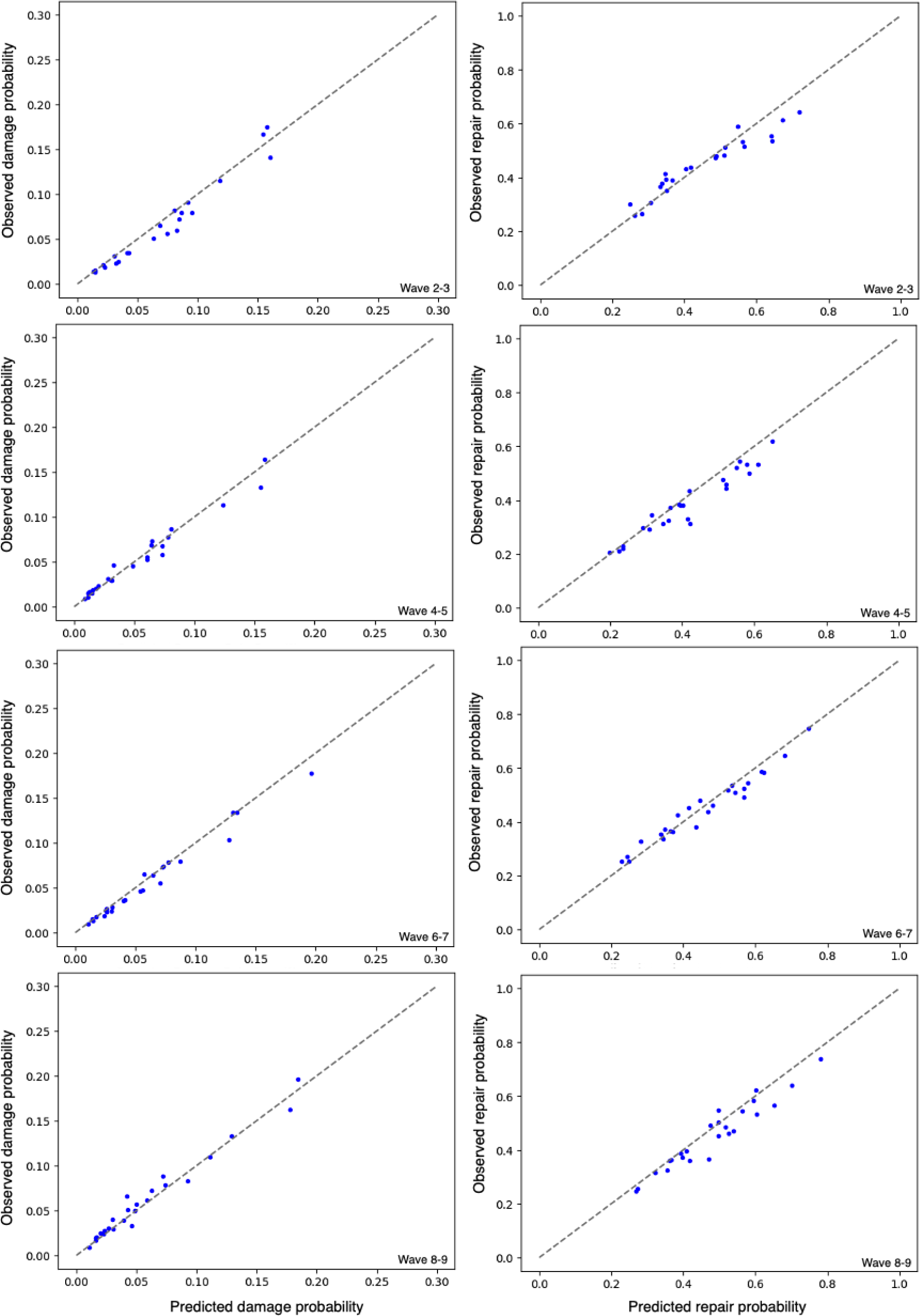
Predicted versus observed average damage and repair probabilities for ADLs. Each point represents one ADL. Waves are as indicated. The average *R*^2^ across waves are in Table III ADLs. The gray dashed line indicates perfect calibration.

**FIG. S4:**
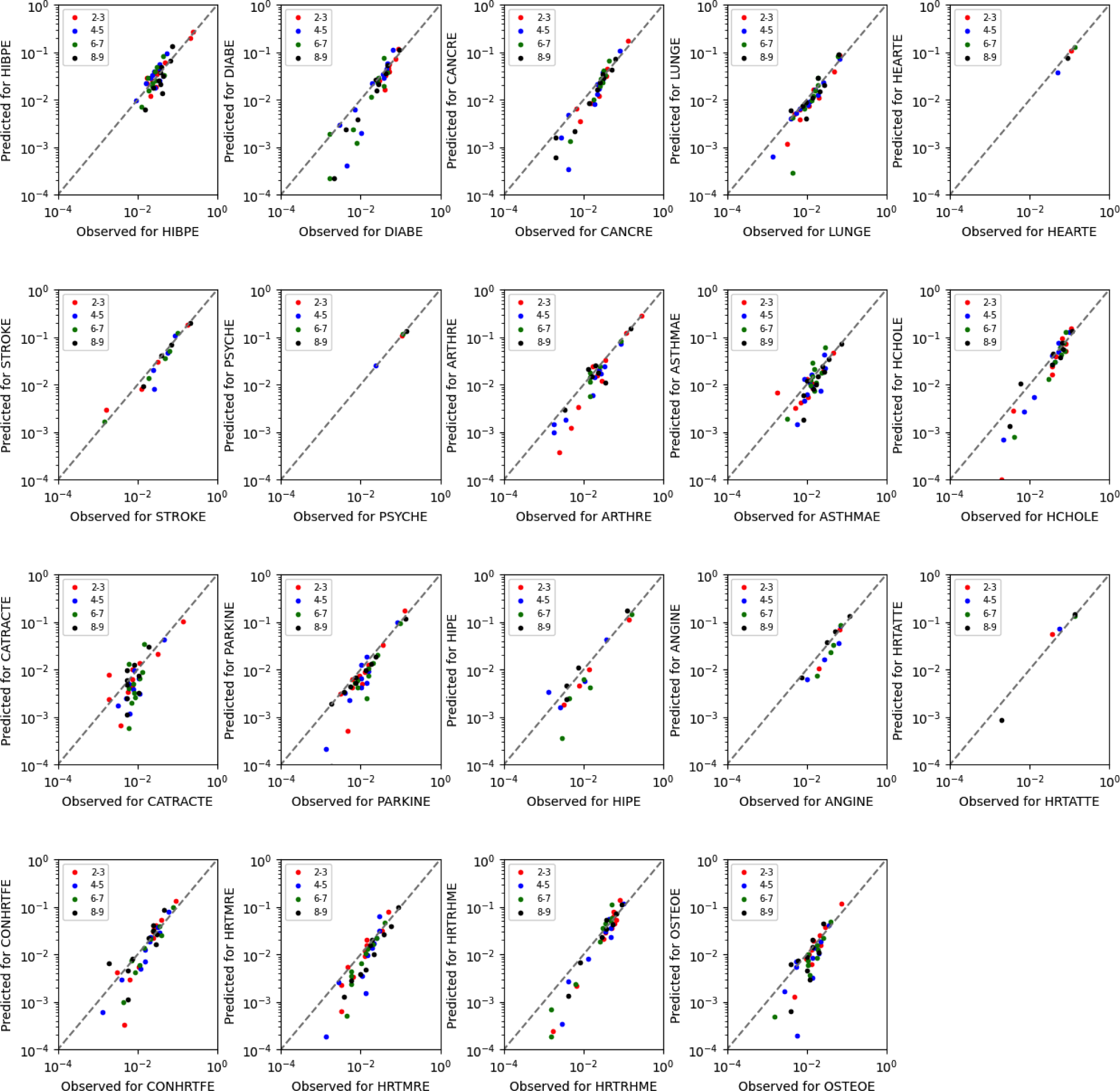
Individual disease calibration curves for damage transition probabilities. Each point is the average of one decile, for the indicated disease, and for the waves indicated by the legend. The gray dashed line indicates perfect calibration.

**FIG. S5:**
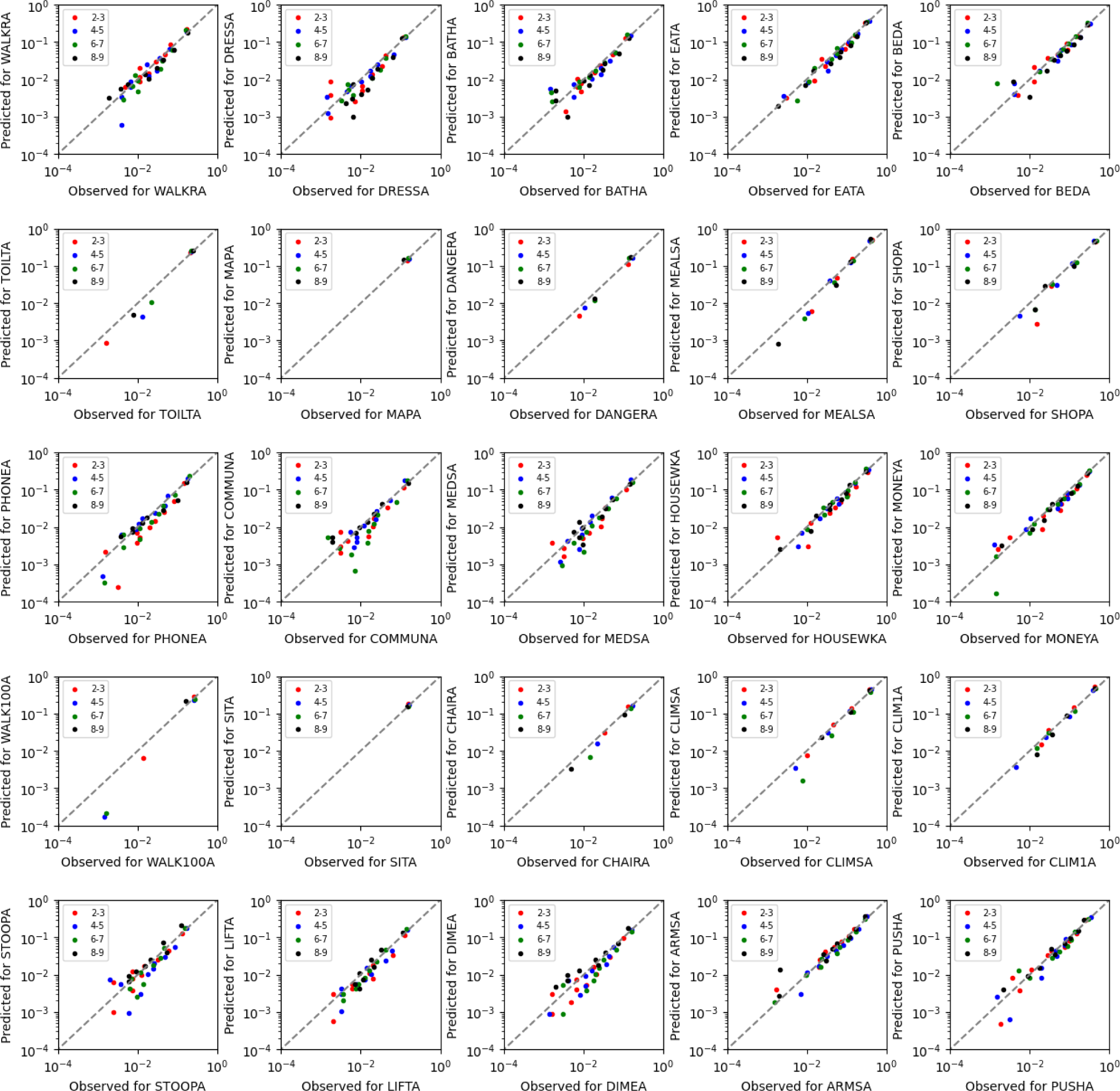
Individual ADL calibration curves for damage transition probabilities. Each point is the average of one decile, for the indicated ADL, and for the waves indicated by the legend. The gray dashed line indicates perfect calibration.

**FIG. S6:**
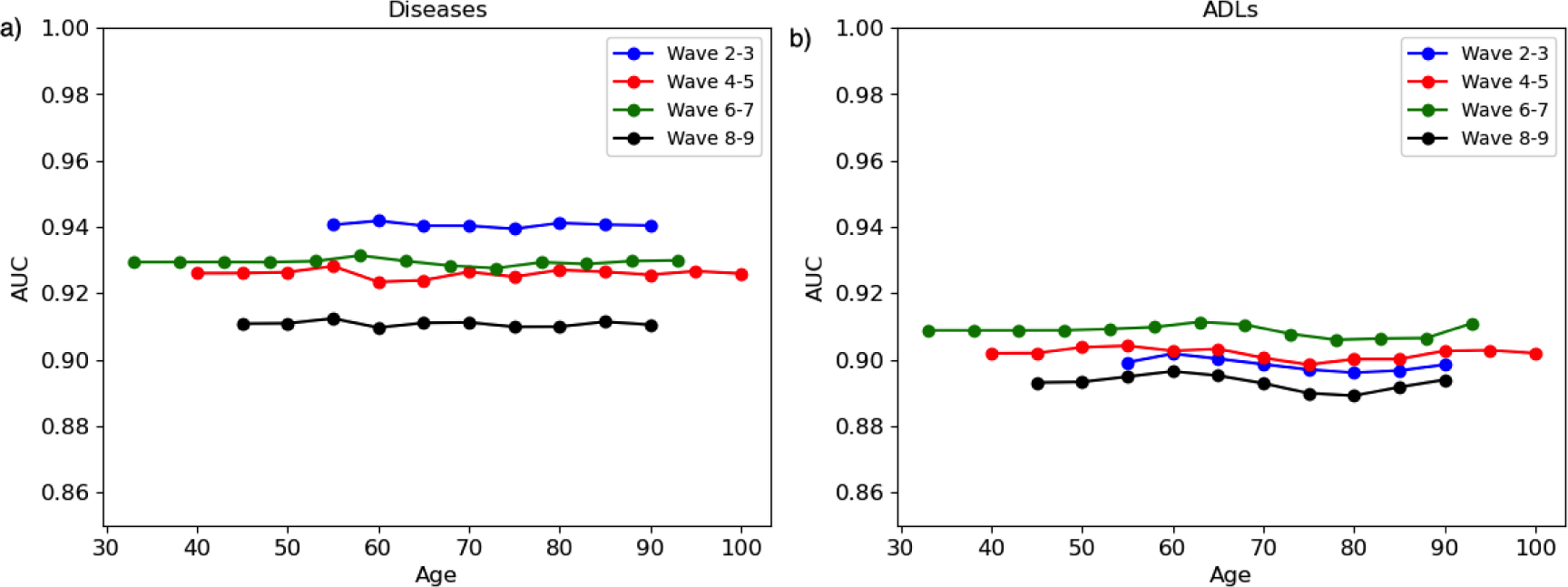
Prediction performance vs age. We show the average AUC of a) disease predictions or b) ADLs for different ages (in five year bins) as indicated, and for different waves as indicated in the legend. Prediction performance does not strongly depend on age.

**FIG. S7:**
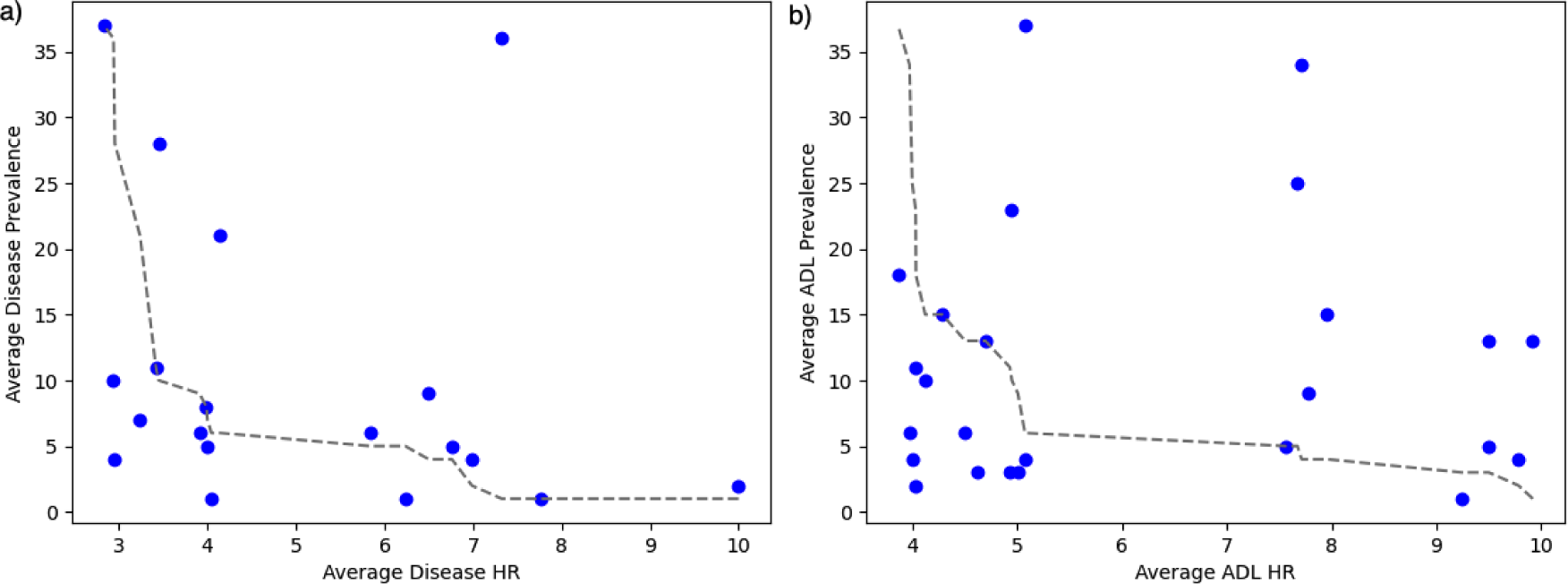
Hazard ratio versus prevalence of a) diseases and b) ADLs. The values are averaged over all wave transitions. The gray dashed curve shows the perfect inverse Spearman’s rank correlation (=1.00) between prevalence and HR values. The measured Spearman’s are −0.43 for diseases and 0.06 for ADLs.

**FIG. S8:**
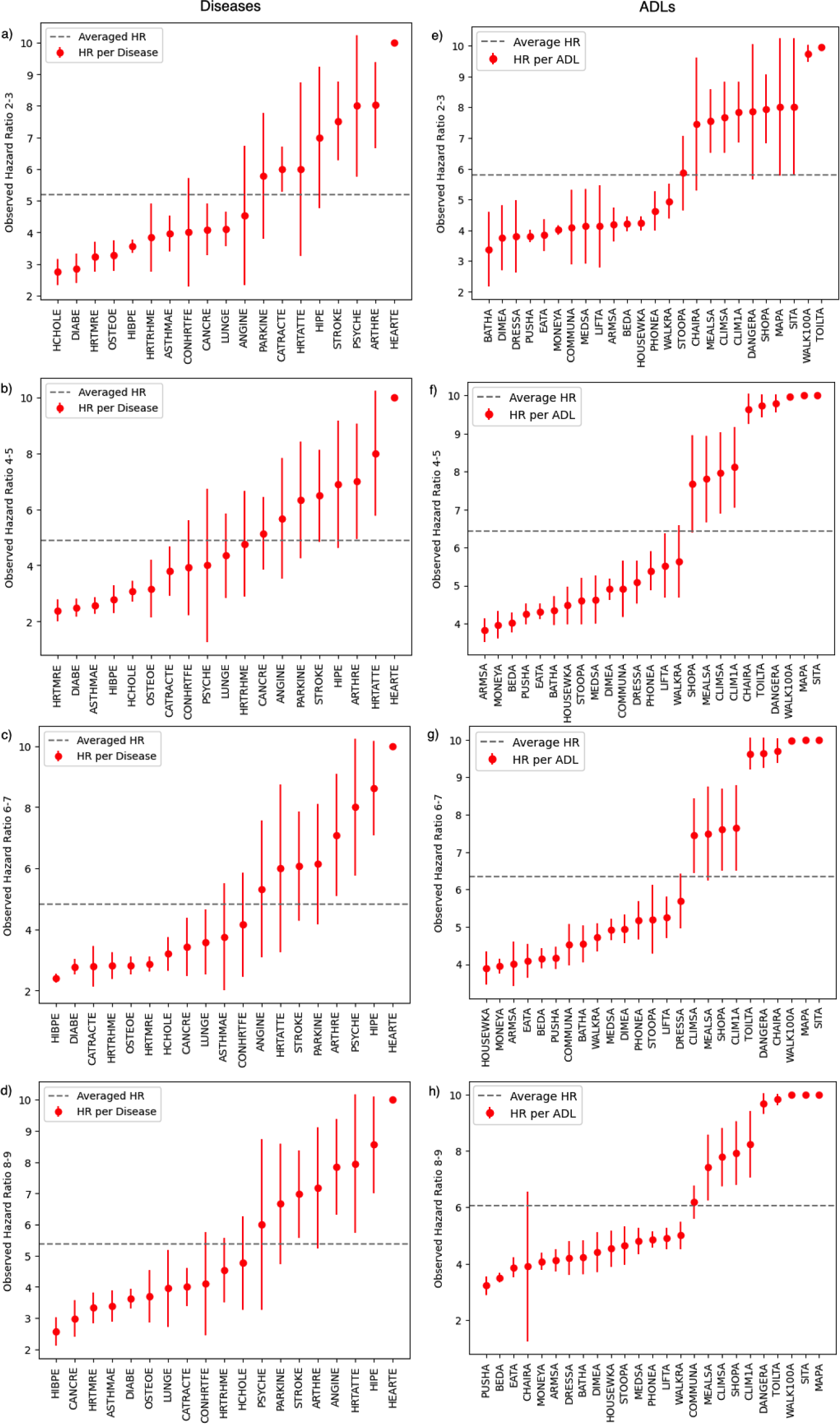
HR versus all diseases (left, a-d) and ADLs (right, e-h) for all wave transitions as indicated. HRs are rank ordered for each wave transition. The horizontal dashed line indicates the average HR over all diseases or ADLs, respectively.

**FIG. S9:**
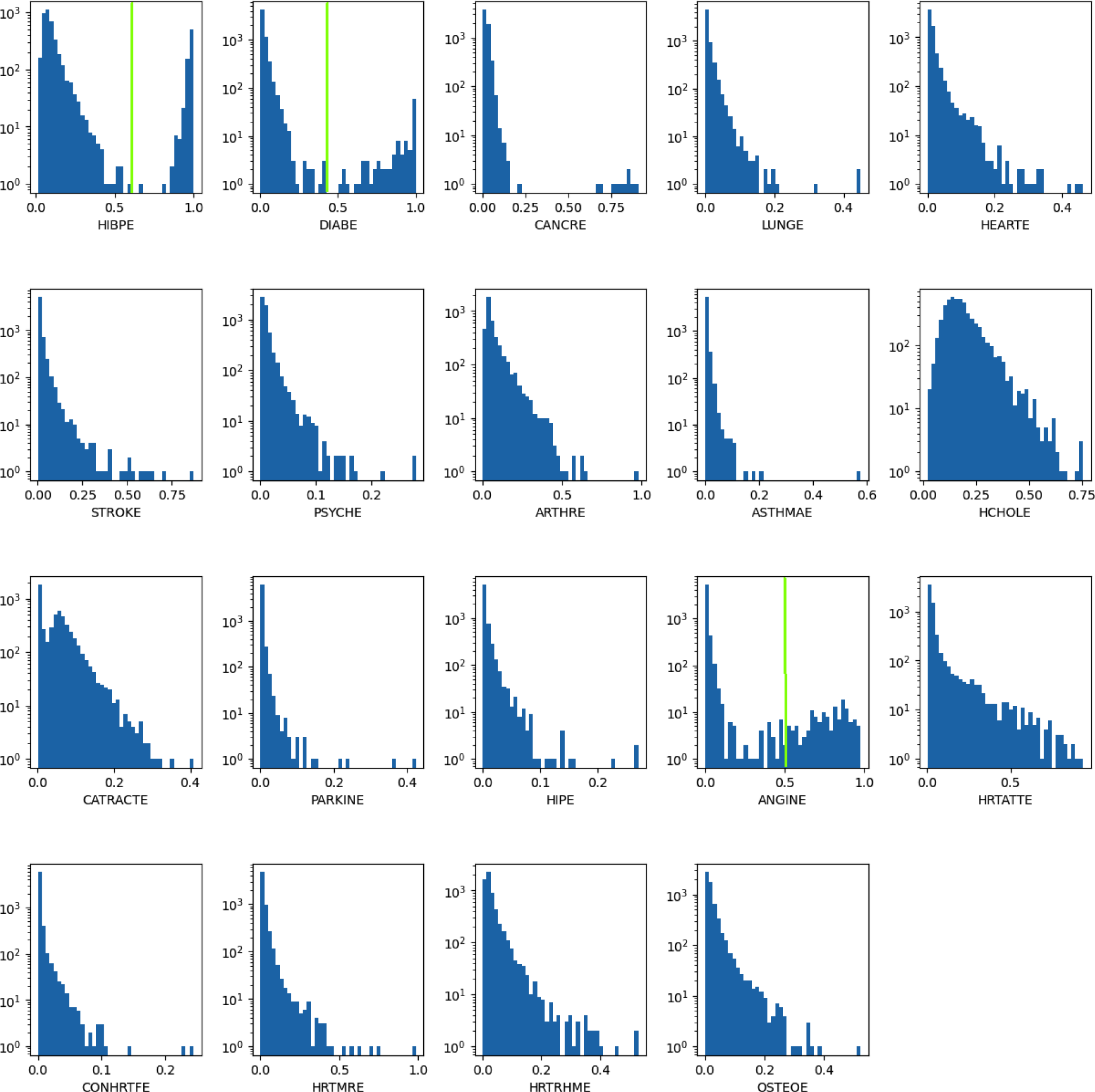
Histograms of predicted disease damage transition probabilities in Waves 2-3. Note the log-scale. All distributions are right-skewed from the mode (peak). The tail appears exponential (straight in this semilog plot) in some distributions (HIBPE, HCHOLE, and CATRACTE) but more generally power law (see Table S10). Some distributions appear bimodal (HIBPE, DIABE, and ANGINE) with vertical green lines indicating the cutpoints used in Table S13.

**FIG. S10:**
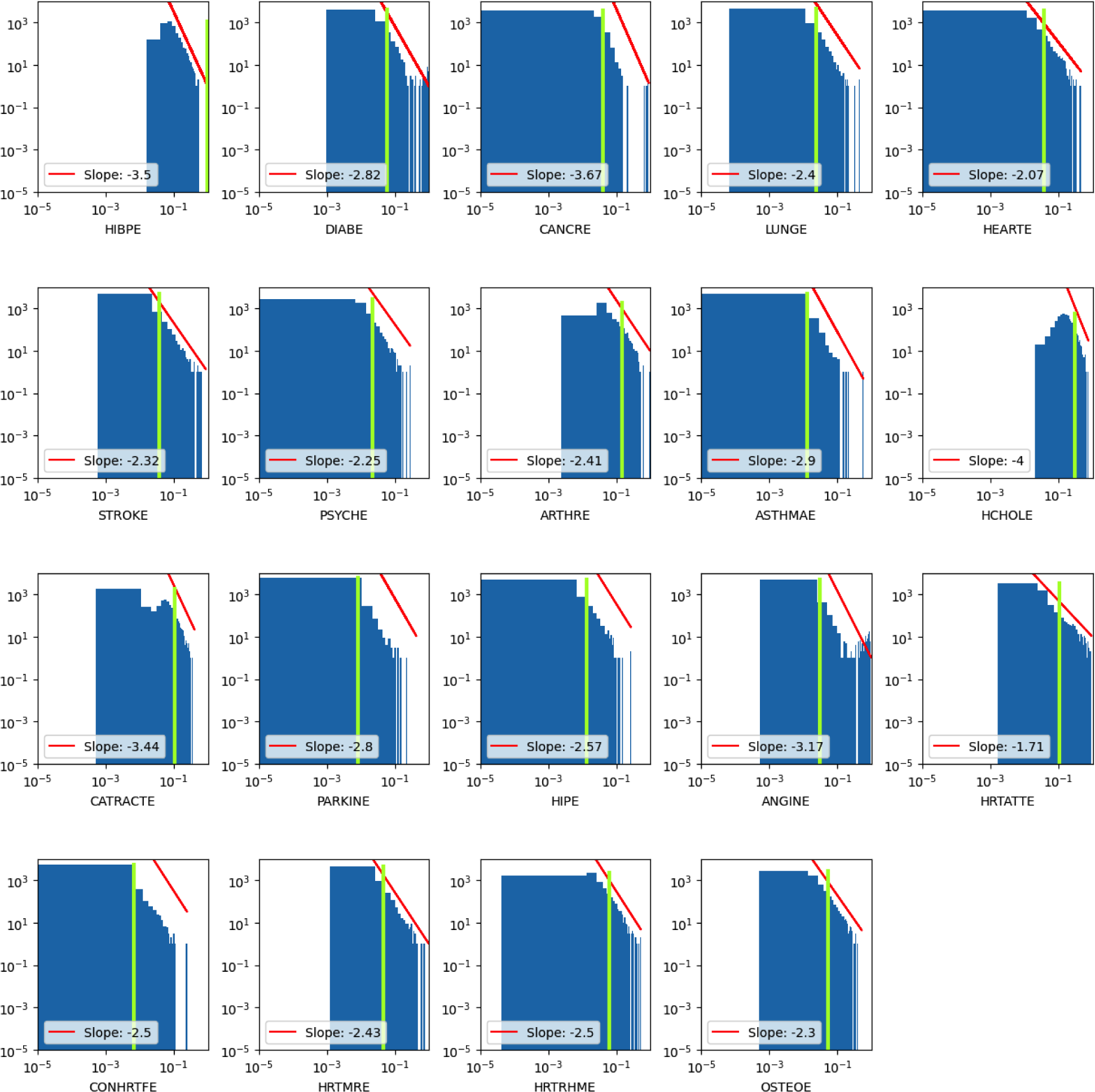
Histograms of predicted disease damage transition probabilities in Waves 2-3. Note the log scale for both axes. Green vertical lines indicate the highest decile of the transition probabilities. The red lines indicate power-law fits by eye, with indicated exponents.

**FIG. S11:**
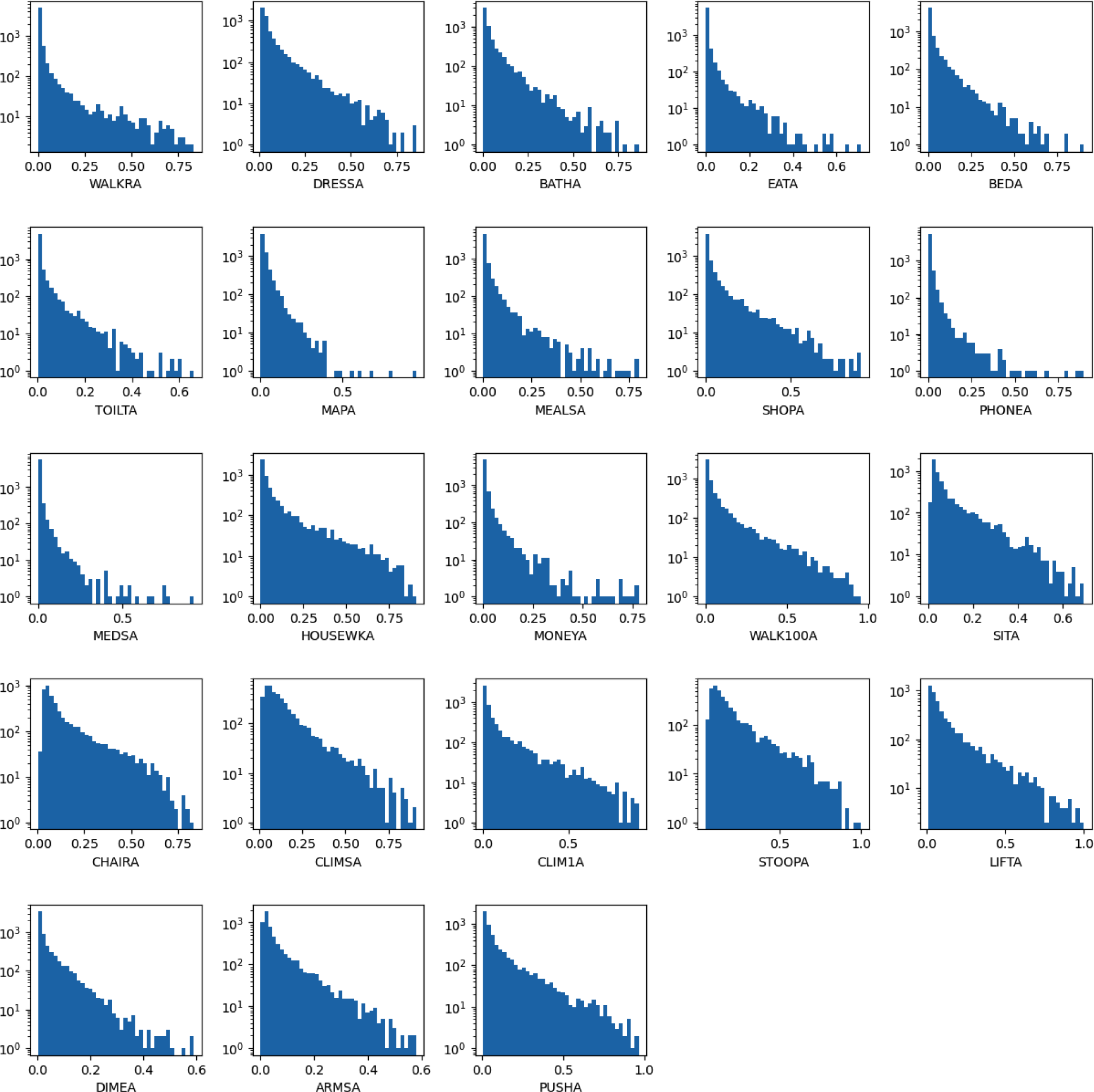
Histograms of predicted ADL damage transition probabilities in Waves 2-3. Note the log-scale. All distributions are right-skewed from the mode (peak). No distributions appear bimodal. All tails appear concave up (i.e. non-exponential).

**FIG. S12:**
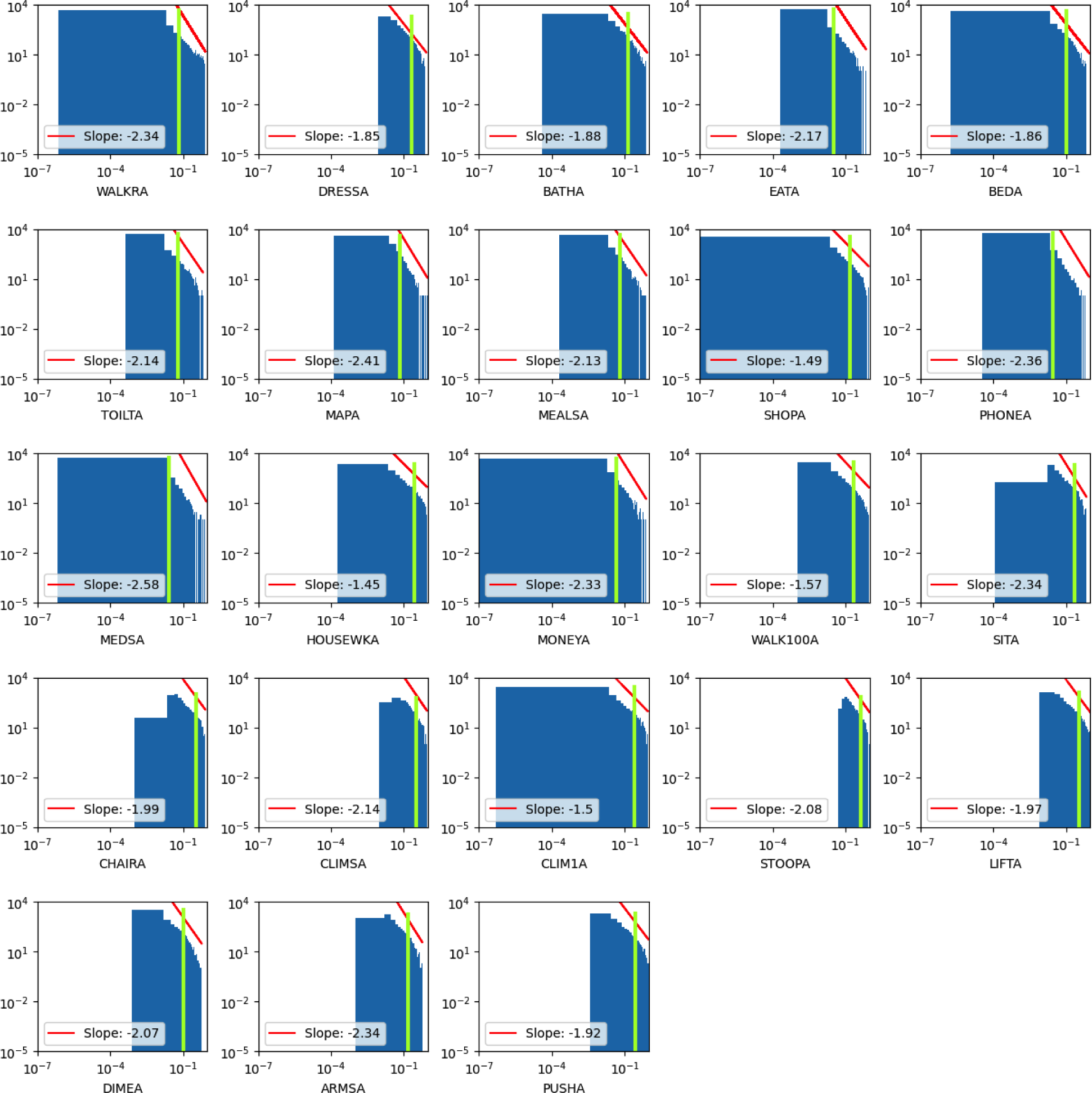
Histograms of predicted ADL damage transition probabilities in Waves 2-3. Note the log scale for both axes. Green vertical lines indicate the highest decile of the transition probabilities. The red lines indicate power-law fits by eye, with indicated exponents.

**FIG. S13:**
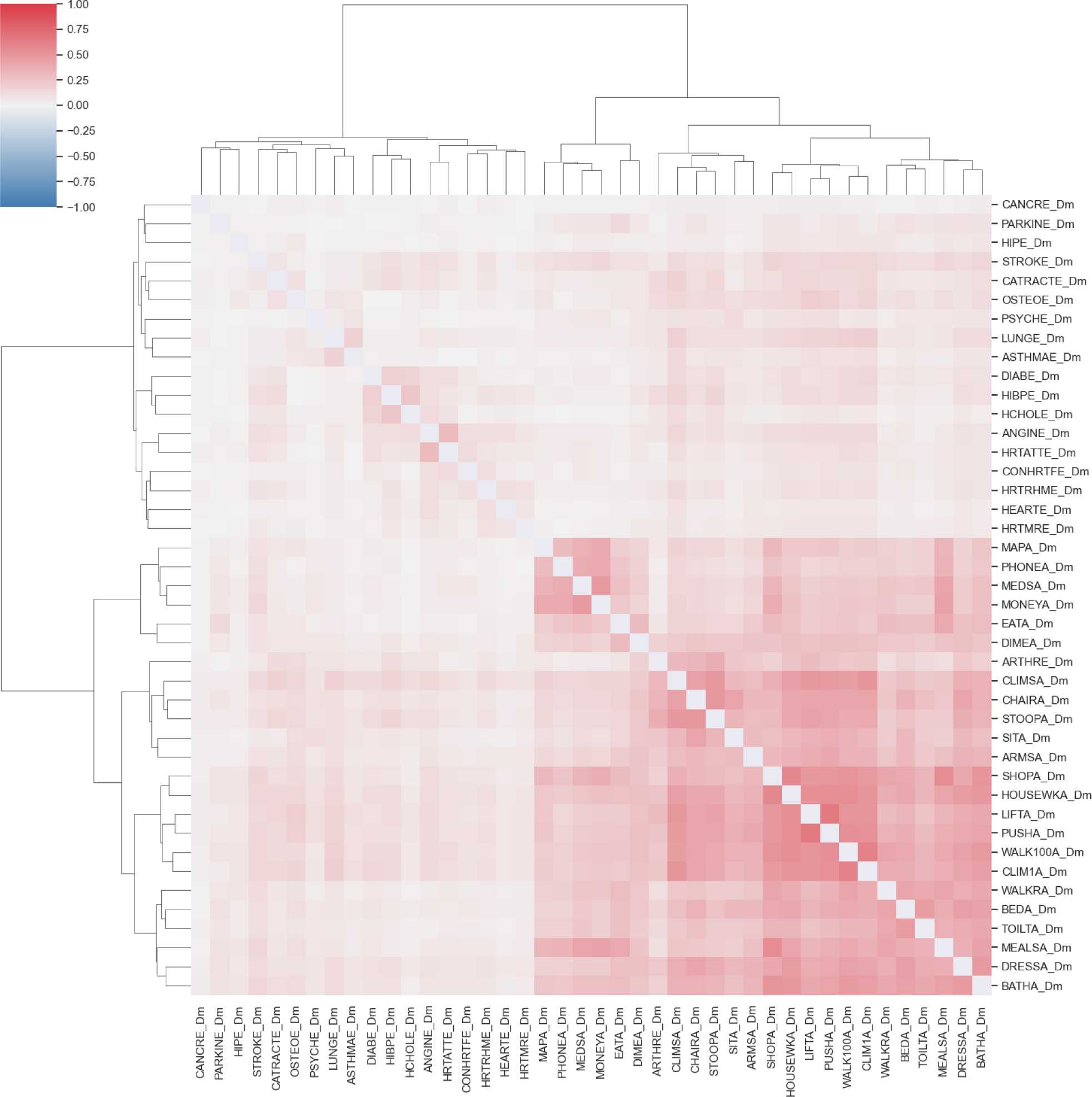
Correlations between observed health states. The heirarchical clustering is also indicated. Note that there is a strong correlation between ADL variables, and a moderate correlation between ADL and disease variables. There is a smaller correlation between disease variables. All of the stronger correlations are positive. We do not show diagonal correlations (i.e. variances).

**FIG. S14:**
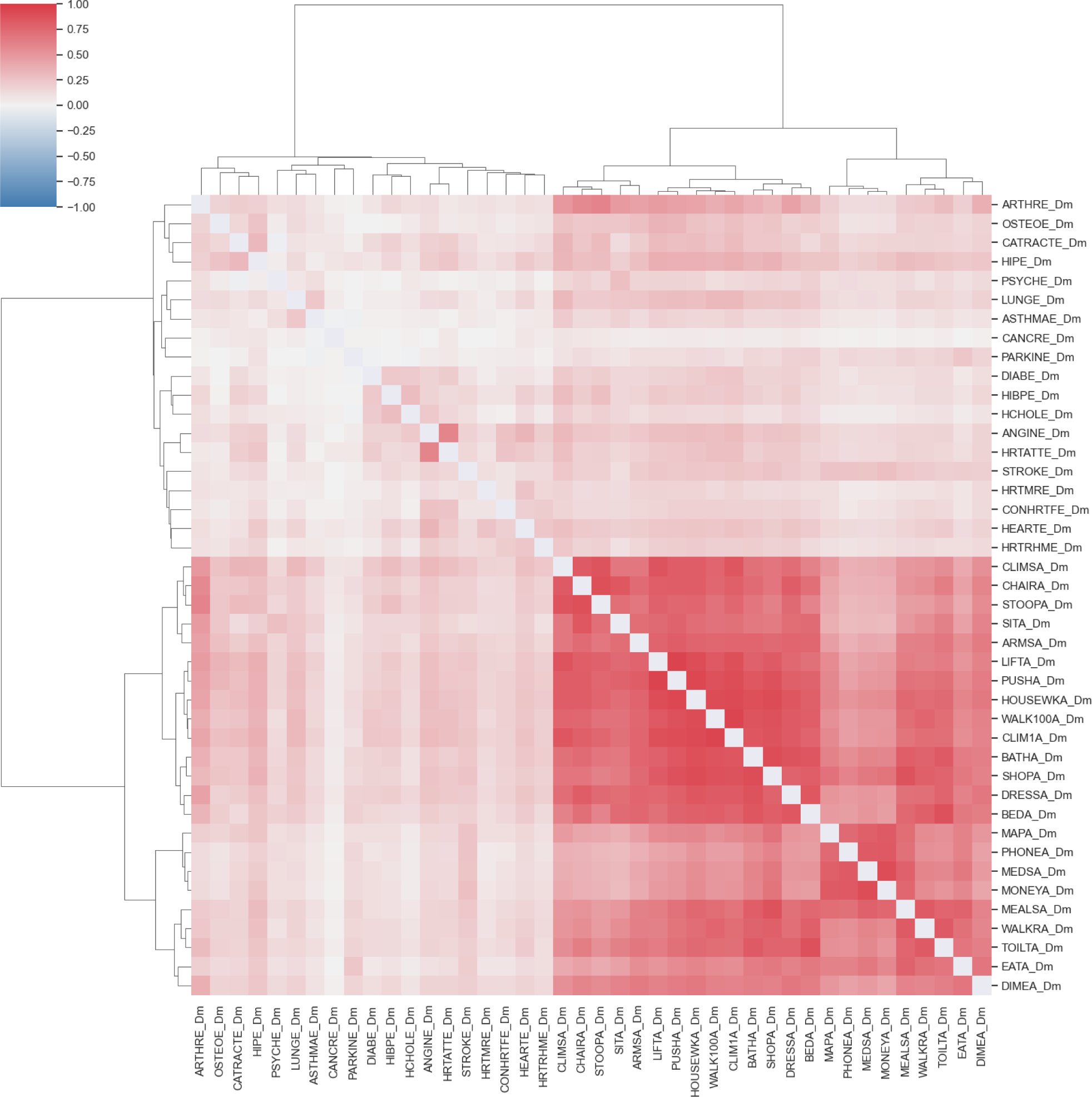
Correlations between predicted health states. The heirarchical clustering and block structure is similar to the observed case in Fig. S13, however the correlations are significantly stronger.

